# Evidence for non-optimal codon choice in highly transcribed sex-biased genes of *Drosophila melanogaster*

**DOI:** 10.1101/2024.11.24.625100

**Authors:** Carrie A. Whittle, Cassandra G. Extavour

## Abstract

Biases in synonymous codon use occur in many unicellular and multicellular organisms. Optimal codons, defined as those most commonly used in highly transcribed genes, are thought to arise from selection for cost-efficient translation, which would favor codons with abundant matching tRNAs. Such presumed selection is described as optimal codon choice. Non-optimal codons, defined as those least commonly used in highly transcribed genes, may in principle also play important roles, but the dynamics of their use remain understudied. Here we examine non-optimal codon use, using sex-biased genes expressed in the gonads of *Drosophila melanogaster* as a case study. We show that genes with sex-biased expression exhibit a preference for non-optimal codon use, especially testis-biased genes. Further we show that the use of non-optimal codons is not random. Instead, specific non-optimal codons are favored, again especially in testis-biased genes. Non-optimal codon use is positively linked to elevated disorder of the encoded proteins. Remarkably, all 18 degenerate amino acids were associated with higher disorder when encoded by the identified primary non-optimal codon, than when encoded by its sister optimal codon. We hypothesize that selection may have promoted non-optimal codon choice for a subset of favored non-optimal codons to regulate translation. We discuss the putative roles of tRNA gene copy numbers, pleiotropy, and sex-biased expression in the evolution of this level of gene regulation.

**Significance Statement:** Optimal codons, those codons most commonly used in highly expressed protein-coding genes, are thought to improve translational efficiency in a range of organisms. However, relatively minimal attention has been given to non-optimal codons, those least often used in highly transcribed genes, and their potential roles in translation. Here, using sex-biased gonadal genes of *Drosophila melanogaster* as a model system, we demonstrate that non-optimal codons are preferentially used in sex-biased genes, particularly within highly expressed testis-biased genes, and are associated with disordered proteins. Moreover, the preference for a specific non-optimal codon (per amino acid) is ubiquitous across all degenerate amino acids. We propose that non-optimal codon use is non-random and may have evolved under selection for roles in translational regulation and protein folding in sex-biased genes, and in a manner associated with their tRNA abundances. The findings have significant implications for the understanding of pacing of translation, protein conformation and protein functionality.

## Introduction

The genetic code contains 18 amino acids encoded by synonymous codons (Ikemura 1981; Sharp et al. 1986; Shields et al. 1988; Liu et al. 2021; Plotkin & Kudla 2011). Optimal codons are defined as those codons most frequently used for amino acids in abundant transcripts, and are thought to promote cost efficient or more accurate translation, a phenomenon often described as optimal codon choice (Ikemura 1981; Hershberg & Petrov 2009). To date, optimal codons have been identified in unicellular (Ikemura 1985, 1981; Akashi 2003) and some multicellular organisms, including insects such as flies, beetles, crickets and mosquitoes (Galtier et al. 2018; Ikemura 1985; Moriyama & Powell 1997; Whittle et al. 2021, 2019; Hambuch & Parsch 2005; Williford & Demuth 2012; Whittle & Extavour 2017; Shields et al. 1988; Behura & Severson 2011). The optimization of codons in highly expressed genes is thought to have resulted from adaptation of the coding sequence (CDS) to favor use of the most common tRNAs in the cells, often measured as the number of matching tRNA genes in the genome, which gives rise to a high ratio of tRNA supply to codon demand (Gingold et al. 2012; Du et al. 2017; Duret 2000; Akashi 2003; Percudani et al. 1997; Zhou et al. 2013; Reis et al. 2004). However, a number of studies have found that some optimal codons have no tRNAs (wobble codons) in certain taxa, a scenario that may slow translation (Whittle et al. 2021, 2019; Stadler & Fire 2011). While studies in multicellular taxa have largely focused on optimal codons (Akashi 2001; Ikemura 1981; Cutter et al. 2006; Ingvarsson 2008; Whittle et al. 2007; Behura & Severson 2011; Duret 2000; Duret & Mouchiroud 1999; Hershberg & Petrov 2009), non-optimal codons, defined as those least commonly used in highly expressed genes, may also have translational roles (Whittle et al. 2021, 2019; Zhou et al. 2015; Fu et al. 2016; Liu et al. 2021; Brule & Grayhack 2017; Tuller et al. 2010; Wang et al. 2011). When non-optimal codons have few or no matching tRNA genes, some data suggest that their roles could include targeted slowing of translation for co-translational protein folding or to prevent ribosome jamming (Yu et al. 2015; Liu et al. 2021; Tuller et al. 2010; Stein & Frydman 2019; Letzring et al. 2010; Pechmann & Frydman 2013; Zhou et al. 2015). In addition, under a somewhat paradoxical scenario, some non-optimal codons have been found to have abundant tRNAs, and yet are not used as optimal codons. In these cases, this high tRNA abundance to codon use ratio could lead to rapid and preferential translation of specific mRNAs, namely, those mRNAs with unusually high use of the non-optimal codons (Whittle et al. 2021, 2019; Guimaraes et al. 2020; Torrent et al. 2018). At present however, studies on non-optimal codon use and its potential molecular consequences remain sparse. Key outstanding questions include whether some non-optimal codons may be favored over others, that is, whether non-optimal codons may exhibit non-random use, and if so, whether favored non-optimal codons may be linked to specific properties of the proteins whose transcripts deploy them.

An attractive model system to study codon use, particularly non-optimal codon use, is sex-biased genes expressed in the gonads of multicellular organisms. Because these genes are thought to play an important role in reproductive success, factors influencing their expression and protein functionality may be especially relevant to fitness. Indeed, pioneering studies in the insect *D. melanogaster* and in certain plants have shown that optimal codon use differs between sexes, and is lower in male-than in female-biased reproductive genes (Hambuch & Parsch 2005; Zhang et al. 2004; Whittle et al. 2007). Thus, gonadal genes exhibit sufficient selection differences in codon use to be detectable on a gene-wide level (e.g., Fop, or frequency of optimal codons (Bahiri-Elitzur & Tuller 2021) (Hambuch & Parsch 2005; Zhang et al. 2004; Whittle et al. 2007)), and thus provide an informative system to fill gaps in our knowledge on the dynamics of non-optimal codon use. *D. melanogaster* as a study system is especially valuable given its vast genomic and gene expression resources available for study (Gramates et al. 2022; Li et al. 2014; Graveley et al. 2011).

Some previous observations suggest that the dynamics of non-optimal codon use in genes may be dependent on protein-folding properties (Liu et al. 2021; Zalucki & Jennings 2007; Yu et al. 2015). For example, studies of the circadian clock proteins FQR (in *Neurospor*a) and PER in *D. melanogaster* have found that experimentally changing non-optimal codons to optimal codons in disordered regions impaired protein form and function, a pattern not found upon optimization of codons in structurally ordered regions (α-helixes) (Zhou et al. 2015; Yu et al. 2015; Fu et al. 2016). Thus, non-optional codons in those two genes may contribute to proper protein structure, particularly for disordered regions (Zhou et al. 2015; Yu et al. 2015). Further, changes in a single synonymous codon, including non-optimal codons, have been linked to altered protein folding and disease phenotypes (Fu et al. 2016; Kimchi-Sarfaty et al. 2007; Letzring et al. 2010; Kirchner et al. 2017; Goodarzi et al. 2016). Thus, the extent and accuracy of protein folding may be a factor linked to a gene’s overall non-optimal codon use (Zhou et al. 2015), or potentially to the use of specific non-optimal codons. One approach to quantify the contribution to protein folding of individual amino acid sites (and thus, on a codon-by-codon level) and also on a gene-wide basis, is to assess the relative solvent accessibility (RSA). The RSA is an index that quantifies the exposed surface area of an amino acid within the protein structure (that reflects secondary and tertiary structures), relative to its maximum possible surface area in Angstroms (Braun 1998; Tien et al. 2013). Low RSA values typically indicate deeply buried hydrophobic amino acids or protein regions, while exposed residues at the protein surface are typically hydrophilic with relatively high RSA values. RSA analyses have proven effective in revealing links between protein folding and the rate of protein sequence evolution, including slow protein evolution in highly ordered (low RSA) regions (Scherrer et al. 2012; Conant & Stadler 2009; Franzosa & Xia 2009; Moutinho et al. 2019; Echave et al. 2016; Bricout et al. 2023; Ramsey et al. 2011). In this regard, it is reasonable to infer that RSA may also provide an effective means to study protein folding properties with respect to optimal and non-optimal codon use.

Here, from the study of highly expressed sex-biased genes in *D. melanogaster*, we provide evidence of non-random non-optimal codon use, that is consistent with selection for the use of non-optimal codons. Specifically, we report a stepwise increase in non-optimal codon use from highly transcribed sexually unbiased, to ovary-biased, to testis-biased genes. We show that certain non-optimal codons are markedly preferred over other sister non-optimal codons, especially in the testis-biased genes. In addition, genes with the highest frequency of non-optimal codon use (>60% of amino acids) show significantly elevated gene wide RSA, and this pattern is most prevalent in testis-biased genes. We further reveal that for a single amino acid, the most strongly favored non-optimal codon per amino acid is consistently linked to elevated RSA as compared to its sister optimal codon, a pattern observed for 18 of 18 degenerate amino acids in the genetic code. The favored use of certain non-optimal codons in testis-biased genes includes both codons with few and with plentiful matching tRNA gene copy numbers. This suggests that selection for specific non-optimal codons in testis-biased genes could enhance at least two potential functions, namely slowed translation and targeted up-translation (Stein & Frydman 2019; Liu et al. 2021; Whittle et al. 2021)). We hypothesize that the use of non-optimal codons in sex-biased genes, and especially in male-biased genes, may be mediated by very low cross-tissue gene pleiotropy (Otto 2004; Mank & Ellegren 2009; Yanai et al. 2005), and by weak or absent potential for sexual conflict (Ellegren & Parsch 2007; Wong & Holman 2023).

## Results and Discussion

### *A Priori* Classifications of Organismal Optimal and Non-optimal codons in *D. melanogaster*

Prior to analyses of sex-biased gene sets and codon use, we first aimed to define the organism-wide optimal and non-optimal codon status for *D. melanogaster* for each of the 59 codons encoding the 18 degenerate amino acids. For this, we compared the relative synonymous codon usage (RSCU) (Sharp & Li 1987) of a set of highly expressed (ribosomal protein genes) versus all genes in the genome, and defined this value as the ΔRSCU_Ribosome-All_ (see Materials and Methods (Subramanian et al. 2022)). This approach allows a means to identify the optimal (most common), and the non-optimal (least common) codons used in highly expressed genes in the genome (as compared to low or moderately expressed genes), which is the foundation for our subsequent analysis of codon use in highly transcribed sex-biased genes (Table S1). All codon classifications obtained with this method, and their tRNA abundance statuses, are provided in Table S1. We identified optimal codons as universally G3 and C3 codons, as previously suggested for *D. melanogaster* (Duret & Mouchiroud 1999; Hambuch & Parsch 2005; Heger & Ponting 2007; Shields et al. 1988). This pattern of codon use is not explainable by mutational biases, but rather likely arose from selection pressures (Shields et al. 1988; Duret & Mouchiroud 1999; Heger & Ponting 2007). Herein, for our study, for each of the 18 degenerate amino acids we explicitly defined the (primary) optimal codon (Whittle et al. 2019, 2021) as that with the largest positive ΔRSCU_Ribosome-All_ value compared with its sister codons (Table S1). All other codons for a given amino acid (a total of 41) were classed as non-optimal codons. These typically had a negative or near-zero ΔRSCU value (and an RSCU value 1.5-fold to six-fold smaller than that of the optimal codon). We then focused on further analysis of these non-optimal codons, which have been minimally considered or excluded in codon use studies to date (Duret & Mouchiroud 1999; Hambuch & Parsch 2005; Shields et al. 1988; Heger & Ponting 2007)(see Supplementary Text File S1 for details).

We classified all optimal and non-optimal codons based on the number of exact matching tRNA genes in the *D. melanogaster* genome (Table S1, GtRNA database (Chan & Lowe 2016)), because tRNA abundances, often well reflected by gene copy numbers, underlie purported translational functions (Gingold et al. 2012; Du et al. 2017; Duret 2000; Akashi 2003; Percudani et al. 1997; Stein & Frydman 2019; Stadler & Fire 2011; Whittle & Extavour 2019)). 12 of the 18 optimal codons had the most abundant perfectly matching tRNA genes among its sister codons (e.g., GCC for Gly with 14 tRNAs; denoted as Opt_high-tRNAs_ status; high/low tRNAs were demarcated by the median of 5 tRNA genes). The other six optimal codons had no matching tRNAs, and were classed as wobble codons (Opt_wobble_; e.g. Pro-CCC). In turn, 26 of the 41 non-optimal codons had low tRNA numbers (Non-opt_low-tRNAs_, status; e.g., ATA for Ile) and the remaining 15 had abundant tRNA genes (Non-opt_high-tRNAs_ status; e.g., ACT for Thr) (Table S1). Hereafter, we focus on the study of sex-biased genes and their patterns of optimal and non-optimal and optimal codon use (Table S1). When pertinent, we also consider statuses of their tRNA gene abundances.

### Identification and Characterization of the Sex-Biased Gene Sets

We identified highly expressed testis-biased (N=916), ovary-biased (N=258) and unbiased (N=605) genes in *D. melanogaster* (Fig. S1A) using the strict criteria that a gene have ≥100 RPKM and at least five-fold-bias in one sex versus the other to be classified as sex-biased (testis versus ovaries (mated)), where genes classified as unbiased had <5 fold bias (and ≥100 RPKM in at least one sex), based on transcriptome data from modEncode (Table S2, (Graveley et al. 2011; Gramates et al. 2022) see Materials and Methods). For stringency, we also compared gene expression in the male accessory glands and virgin ovaries (Table S2). We found that 86.9% of the identified testis-biased genes were also male-biased in accessory glands (using a cutoff of ≥ 5-fold sex-bias), and that 77.1% of the identified ovary-biased genes were also female-biased in virgin ovaries, indicating that strong male-female transcriptional differences were also present in those alternate reproductive tissue types. The predicted functions of these genes based on GO analysis showed testis-biased genes were strongly linked to testicular roles such as spermatogenesis, and ovary-biased genes were associated with female reproductive process roles such as the chorion and oogenesis. In contrast, unbiased genes were preferentially involved in the ribosome and other housekeeping functions (Table S3; chromosomal locations of genes are discussed in Supplementary Text File S1). In terms of characteristics of these genes with sex-biased expression, the CDS lengths were shorter for testis-biased (median=273 codons) and unbiased genes (median=269 codons) than for the ovary biased genes (median=498 codons; MWU tests P<0.05) (Fig. S1B). We then determined the *tau* value, an index of gene expression specificity with values from 0 to 1 (Yanai et al. 2005), across 59 tissue stages in *D. melanogaster* (Table S2). *tau* exhibited a stepwise decline from testis-biased genes (median=0.95) to ovary-biased genes (median=0.83) to unbiased genes (median=0.74) (ranked ANOVA and Dunn’s P<0.05 for each paired contrast; median genome-wide *tau=*0.85, Fig. S1C). Thus, the testis-biased genes exhibited higher specificity, and lower cross-tissue pleiotropy (Meisel 2011; Assis et al. 2012; Mank & Ellegren 2009), than ovary-biased and unbiased genes.

We assessed the relationships between tRNA gene numbers, amino acid use, and amino acid size complexity (S/C) scores, which are thought to reflect biosynthetic costs of amino acids (provided in Table S4 (Dufton 1997). The tRNA gene copy number per amino acid (pooled across all codons) was strongly positively correlated to the amino acid use (percentage of each amino acid relative to all concatenated amino acids per gene set) for all genes (testis-biased genes: Spearman’s R=0.91, P<10^−7^; ovary-biased genes: R=0.85, P<10^−7^; unbiased genes: R=0.83, P<10^−7^; Fig. S2A-C). Moreover, the S/C score per amino acid (Dufton 1997) was inversely correlated to the organismal tRNA gene numbers (R=-0.59, P<10^−7^, Fig. S2D). In sum, the patterns suggest that all three gene sets have a history of selection on codon use (Du et al. 2017; Akashi 2003; Whittle et al. 2021), further affirming their utility as a model to study dynamics of codon use in this taxon.

By examining the codon adaptation index (CAI), an index that reflects the gene-wide codon frequencies relative to a highly expressed reference gene set (here, ribosomal genes) (Subramanian et al. 2022; Sharp & Li 1987), we found a stepwise decrease in CAI from unbiased, to ovary-biased, to testis-biased genes (ranked ANOVA and Dunn’s P<0.05, Fig. S3A; note that CAI was highly correlated to Fop across the genome, Spearman’s R=0.967, P<10^−7^). In other words, as genes exhibited increased gonad-biased expression, they also presented increased non-optimal codon use, with the highest non-optimal codon use in testis-biased genes (see Supplementary Text File S1). Further, within our highly expressed sex-biased genes sets (the targeted gene sets herein), we subdivided expression into two classes, ≥200 RPKM (extremely high expression), and ≥100-200 RPKM (high expression) and subdivided genes by CDS length (short and long CDS lengths, with long defined as ≥272 codons; cutoff at the 33^rd^ percentile of all gene lengths), which were linked to CAI within each of the sex-biased gene sets (Fig. S3B). The differences in CAI between gene sets (Fig. S3A) were robust to both transcription level and length cutoffs (Fig. S3A, B; see also Supplementary Text File S1). The high non-optimal codon use in testis-biased genes is not due to enhanced genetic interference, as those processes would be expected to ameliorate non-optimal codon use in longer rather than shorter genes, a pattern that was absent from our data (Fig. S3) (Loewe & Charlesworth 2007; Betancourt & Presgraves 2002; Comeron et al. 1999; Hambuch & Parsch 2005). The testis-biased genes herein were short (Fig. S1B), and even within the testis-biased gene set (Fig. S3B), shorter genes (<272 codons) had greater non-optimal codon use (lower CAI) than longer genes for both expression classes ((MWU-tests P<0.05, Fig. S3B; see Supplementary Text File S1). Thus, other factors must explain the stepwise increase in non-optimal codon use from unbiased, to ovary-biased to testis-biased genes. Moreover, it is notable that the extremely highly expressed testis-biased genes tended to have an elevated degree of sex-bias relative to the highly expressed testis-biased genes (the median was 149.5 and 210 fold bias for long and short genes in the former, and 71.0 and 69.5 fold bias in the latter, consistent with high extent of specialization in the testis in each group), suggesting the more biased the expression in the testis (among genes with ≥100RPKM), the greater the use of non-optimal codons.

### Non-optimal Codon Use is Non-random in *D. melanogaster* Sex-biased Genes

To further characterize codon use in unbiased and sex-biased genes, we assessed codon use on a codon-by-codon basis for each gene set described above. We determined the ΔRSCU between testis-biased genes versus ovary-biased genes (ΔRSCU_Testis-Ovary_), and between testis-biased genes versus unbiased genes (ΔRSCU_Testis-Unbiased_), for each of the 59 codons with synonymous codons (*cf.* (Whittle et al. 2007)). In these contrasts, a positive ΔRSCU value indicates higher use of a codon in the testis-biased genes, and a negative value denotes a reduction in its use within testis-biased genes (and thus elevated use in the respective ovary or unbiased genes, see Materials and Methods). The ΔRSCU_Testis-Ovary_ and ΔRSCU_Testis-Unbiased_ values per codon are shown in Table S5. Remarkably, we found that testis-biased genes reduced their use of optimal codons, and this pattern was universal to all of the degenerate amino acids. Specifically, 16 of the 18 optimal codons in *D. melanogaster* had a statistically significant negative ΔRSCU_Testis-Ovary_ (P<0.05, only exceptions were for Ser and Thr P>0.05), and 18 of 18 optimal codons had a negative value for ΔRSCU_Testis-Unbiased_. The magnitude of negative ΔRSCU values was consistently larger for the ΔRSCU_Testis-Unbiased_ contrast than for the ΔRSCU_Testis-Ovary_ contrast, consistent with a stepwise decline in the use of optimal codons from unbiased genes to ovary-biased genes to testis-biased genes (Table S5, Fig. S3). This suggests the possibility that there are weaker translational selection pressures for optimal codons in both types of sex-biased genes, and especially in the testis-biased genes. However, another significant hypothesis worthy of consideration here is that the patterns may reflect a non-random displacement of optimal codons by the favored use of non-optimal codons in testis-biased genes. In other words, it could be the case that testis-biased genes reduce their use of optimal codons not arbitrarily, but rather in favor of preferential use of non-optimal codons.

To test this hypothesis, we asked whether there was any pattern to the non-optimal codons that were used in testis-biased genes. We found that for the nine amino acids encoded by two synonymous codons (Asn, Asp, Cys, Gln, Glu, His. Lys, Phe, Tyr), the universal reduction in use of the optimal codon in testis-biased genes was compensated by increased use of the sister non-optimal codon (as indicated by positive ΔRSCU_Testis-Unbiased_ and ΔRSCU_Testis-Ovary_ values, Table S5), consistent with a hypothesis of preferential use of the non-optimal codons in testis-biased genes. Nonetheless, the two-fold degenerate amino acids cannot be used to fully differentiate whether there may also be codon choice among the non-optimal codons. Thus, we examined the nine amino acids that were encoded by three, four or six synonymous codons (Ala, Arg, Gly, Ile, Leu, Pro, Ser, Thr, Val). If the low use of optimal codons in testis-biased genes were solely due to weaker selection for optimal codons, then one would expect a similar degree of enhanced use of all the synonymous non-optimal codons per amino acid in testis-biased genes (under a neutral model). In contrast, however, we found that the elevated use of non-optimal codons in testis-biased genes was non-arbitrary (Fig. 1A, Table S5). As an example, for the four-fold degenerate amino acid Thr, a neutral model would predict similar and increased use of the three non-optimal codons (ACG, ACA, ACT) under reduced use of the optimal codon (ACC). However, we found that only ACT (Non-opt_high-tRNAs_) had elevated use in testis-biased genes versus ovary-biased genes (ΔRSCU_Testis-Ovary_ =+0.166; t-test P<0.05. Fig. 1B), while the other two non-optimal codons showed no elevation in use. This is not consistent with a neutral model of non-optimal codon use, and instead suggests a preference for one specific non-optimal codon over the others (Fig. 1B). The same preference for ACT was also observed using the ΔRSCU_Testis-Unbiased_ comparison (ΔRSCU=+0.201, Table S5), a codon that was detected even though the ovary-biased and unbiased genes were from fully non-overlapping gene sets, indicating that the preference for this non-optimal codon in the testis-biased genes is robust to the two contrasts. Another example is Ile (three-fold degenerate) where the reduced frequency of the optimal codon ATC in testis-biased versus unbiased genes contrast was accompanied by a strong preferential use of just one of the two non-optimal codons (ATA; ΔRSCU_Testis-Ovary_ =+0.114; 9.5 fold higher than that of its synonymous non-optimal codon ATT; Table S5), and the same non-optimal codon preference was also identified using the ΔRSCU_Testis-Unbiased_ contrast (value was +0.266. t-test P<0.05, Fig. 1C, Table S5). While the same primary non-optimal codons (largest positive ΔRSCU) tended to occur in both the ΔRSCU_Testis-Ovary_ and the ΔRSCU_Testis-Unbiased_ contrasts, the latter were more often statistically significant (Fig. 1, Table S5). This likely reflects not only the larger ΔRSCU values in the latter contrasts, but also the larger N values and thus power for the unbiased gene set (N=605 for unbiased, and 258 for ovary-biased). Overall, the pattern of the highly favored use of a specific non-optimal codon over others per synonymous codon family in testis-biased genes was also observed for Gly (GGA, Fig. 1D), Ala (GCA), Leu (TTG), Ser (AGT) and Val (GTA as primary non-optimal codon, GTT as secondary), Pro (CCT, particularly ΔRSCU_Testis-Ovary_ contrast) and Arg (AGG as primary, AGA as a close secondary) (Table S5, Fig. 1A). All primary non-optimal codons ended in A3, T3 or G3 (Table S5), thus excluding a role of mutational biases for a particular nucleotide at the 3^rd^ “silent” position, and suggesting instead that selective processes have influenced the use of those codons. Notably, the identified primary non-optimal codon per amino acid always had a positive value for both ΔRSCU_Testis-Ovary_ and for ΔRSCU_Testis-Unbiased_, (P<0.001 for all latter contrasts) but was typically smaller in value for the former contrast, as expected if non-optimal codons are preferred in sex-biased over unbiased genes (and also more favored in testis-biased than ovary-biased genes). Together, these patterns show that testis-biased genes show lowered use of use optimal codons, and preferentially use specific non-optimal codons, rather than randomly choosing among the available non-optimal codons.

**Fig. 1.**
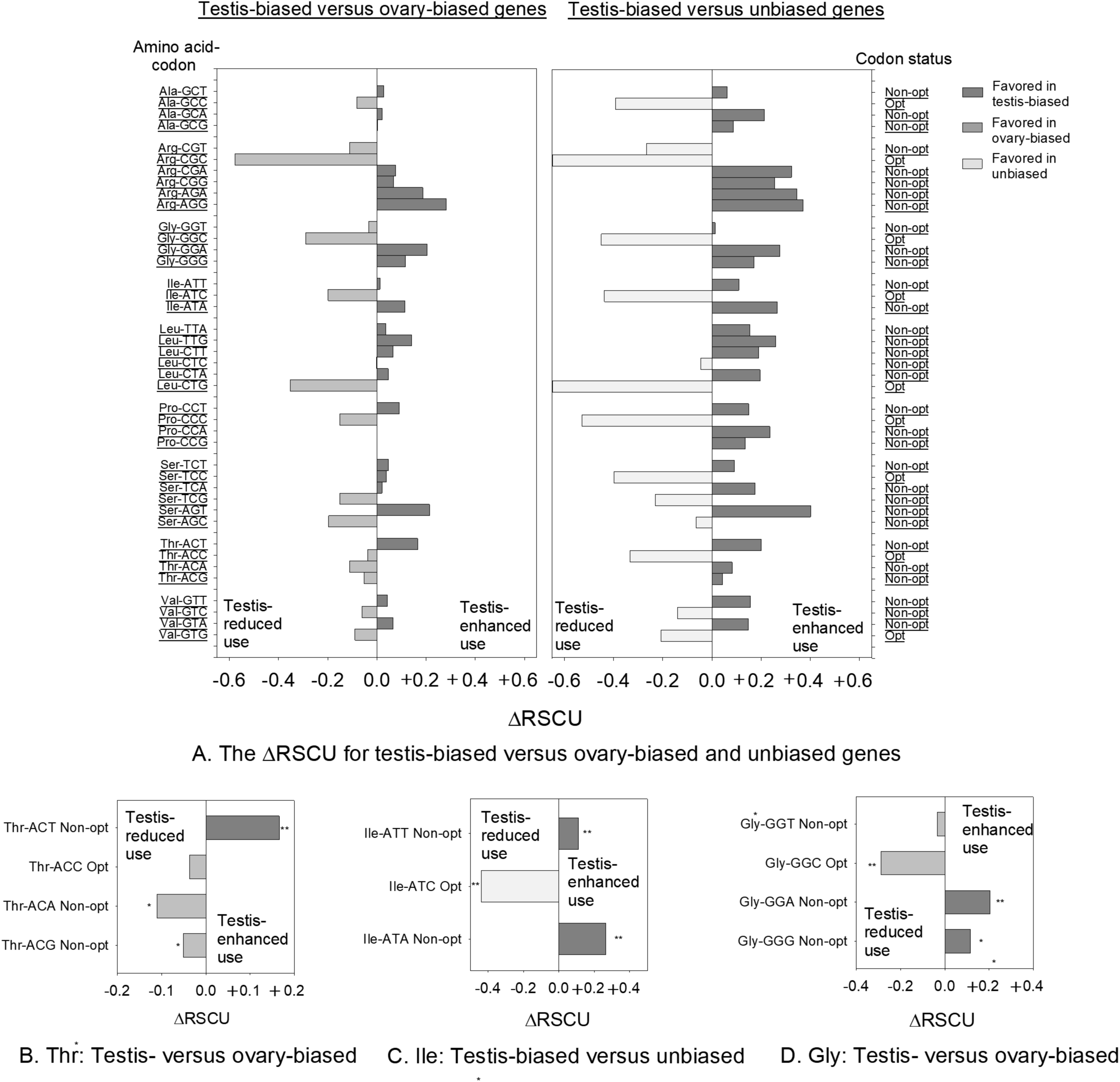
A) The values for ΔRSCU_Testis-Ovary_ (left panel) and ΔRSCU_Testis-Unbiased_ (right panel) per codon for amino acids with three or more synonymous codons (P-values in Table S5). The amino acid-codon and the codon status (left) with respect to tRNAs (right) are each indicated. Close-up examples of the ΔRSCU for B) all four codons for Thr; C) all three codons for Ile; and D) all four codons for Gly (* indicates P<0.05 in B-D). In panel A, ΔRSCU_Testis-Unbiased_ for Leu-CTG Opt_high-tRNAs_ =0.818 and Arg-CGC Opt_wobble_=1.11 and were outside the shown limit of X-axis limit.

Notably, the primary non-optimal codon per amino acid (Table S5, Fig. 1A) in testis-biased genes was not predicted by tRNA status. Specifically, for Arg, Leu and Ser, the primary non-optimal codon had low tRNA abundance (Non-opt_low-tRNAs_ status), while others such as Gly, Pro and Thr favored the use of codons with highly abundant tRNAs (Non-opt_high-tRNAs_ status (Table S5)). We speculate that this pattern suggests that selection of non-optimal codons with few tRNAs (Arg, Leu and Ser) may typically be involved in slowing translation for protein folding or to limit ribosome jamming, while those with abundant tRNAs (Gly, Pro and Thr) may often be involved in ensuring up-translation of mRNAs carrying those codons. These predictions are based on presumed supply-demand ratios of tRNAs:codons (Brackley et al. 2011; Zhou et al. 2015; Stein & Frydman 2019; Brule & Grayhack 2017; Whittle et al. 2021). Thus, we speculate that the primary non-optimal codons identified here could regulate gene function by influencing translation. It should be noted that the tRNA populations for *D. melanogaster* closely match those of its sister species *D. simulans, D. sechellia*, *D. yakuba* and *D. erecta* (Spearman’s R≥0.998 (P<10^−7^), and thus these tRNA populations have existed for a substantial period of time (at least 13 Mya since divergence (Tamura et al. 2004) (see Supplementary Text File 1 for details).

We determined the percentage of genes in each gene set under study that exhibited extreme use of the primary non-optimal codons (defined as RSCU values ≥1.5). As shown in Fig. S4A, a larger percentage of testis-biased genes exhibited extreme use of the primary non-optimal codons than did the comparable ovary-biased and/or unbiased genes (Chi-square P<0.05; shown as examples are Thr-ACT, Gly-GGA, Ile-ATA and Val-GTA; see Supplementary Text File S1 for parallel data for optimal codons, Fig. S4B). It is worth noting that the percent of genes with extreme use of optimal codons with plentiful tRNAs differed from those optimal codons with wobble status (Fig. S4B; in all gene sets, P<0.05). This observation is important as it allows us to infer that these two classes of optimal codons may not be equivalent in functions (see Supplementary Text File S1), and thus in future studies should not be treated as equal optimal codons, nor studied without including tRNA gene data.

### Amino Acids Using Non-optimal Codons Have High Surface Exposure and Disorder

To determine whether these patterns of non-optimal codon use were linked to specific biophysical properties of the encoded proteins, we first assessed the solvent accessibility RSA value for every amino acid (and its associated codon) in the testis-biased genes (N=299,817 amino acids), ovary-biased genes (155,282 amino acids) and unbiased genes (218,580 amino acids). We found that RSA value was strongly correlated to the disorder probability scores (from NetsurfP (Hoie et al. 2022)) in all three datasets (Spearman’s R ≥0.68 for each of three datasets, P<10^−7^). Thus, higher RSA reflects both surface exposure and intrinsic disorder (Piovesan et al. 2022; Akdel et al. 2022; Ilzhofer et al. 2022). In terms of major secondary structures of proteins (α-helices, β-strands and random coils (Hoie et al. 2022)), a majority of amino acids in the proteins of all three gene sets were located within random coils (60.6% for testis-biased, 57.2% of unbiased, and 70.8% for ovary-biased genes, Chi-square tests P<0.05). α-helices were the next most common structures in all three gene sets (31.1, 22.4 and 31.5%, respectively), and β-strands the least common (11.2, 8.3 and 6.8%, Fig. S5A). The lowest RSA values occurred for amino acids located in β-strands, while these values were intermediate for α-helices and highest for random coils for all three gene sets (Fig. S5B, median RSA values across all residues for testis-biased genes were 0.13, 0.34 and 0.53, for ovary-biased genes were 0.11, 0.33 and 0.60, and for unbiased genes were 0.11, 0.30 and 0.57 for β-strands, α-helices and random coils respectively; MWU-tests P<0.05 for all contrasts per gene set). These patterns are consistent with disordered regions having the greatest surface exposure.

For each gene under study, we then determined the percent of amino acids encoded by non-optimal codons, as a gene-wide direct measure of non-optimal codon use (denoted as Percent-Non-opt). The genes were placed into bins of low (≤50%), moderate (>50 to ≤60%) and high Percent-Non-opt codon use (>60%; note that we excluded Met and Typ from this analysis, discussed in Supplementary Text File S1). We found the average RSA per gene increased from the lowest Percent-Non-opt bin (median RSA=0.40), to the moderate (median=0.43) to high (median=0.47) bins (MWU-tests each P<0.05; note the variation within each group within the box plots) in testis-biased genes (Fig. 2A). In other words, the elevated non-optimal codon use in these testis-biased genes tended to be linked to greater disorder in their encoded proteins, whereas genes using more optimal codons tended to be more well-folded (Fig. 2A). For ovary-biased genes and unbiased genes, no statistically significant differences were observed in RSA between the highest Percent-Non-opt bin and the moderate and low bins (P>0.05, Fig. 2B, C), likely because the highest non-optimal codon class was so uncommon in those gene sets for testing (N≤23 genes in each case). Nonetheless, the moderate Percent-Non-Opt bins had elevated average RSA per gene as compared to the low bins for both the ovary-biased and the unbiased genes (MWU-tests P<0.05). Overall, elevated non-optimal codon use is associated with greater protein surface exposure in testis-biased, ovary-biased and unbiased genes, with the most striking stepwise correlation observed for the testis-biased genes (Fig. 2A-C).

**Fig. 2.**
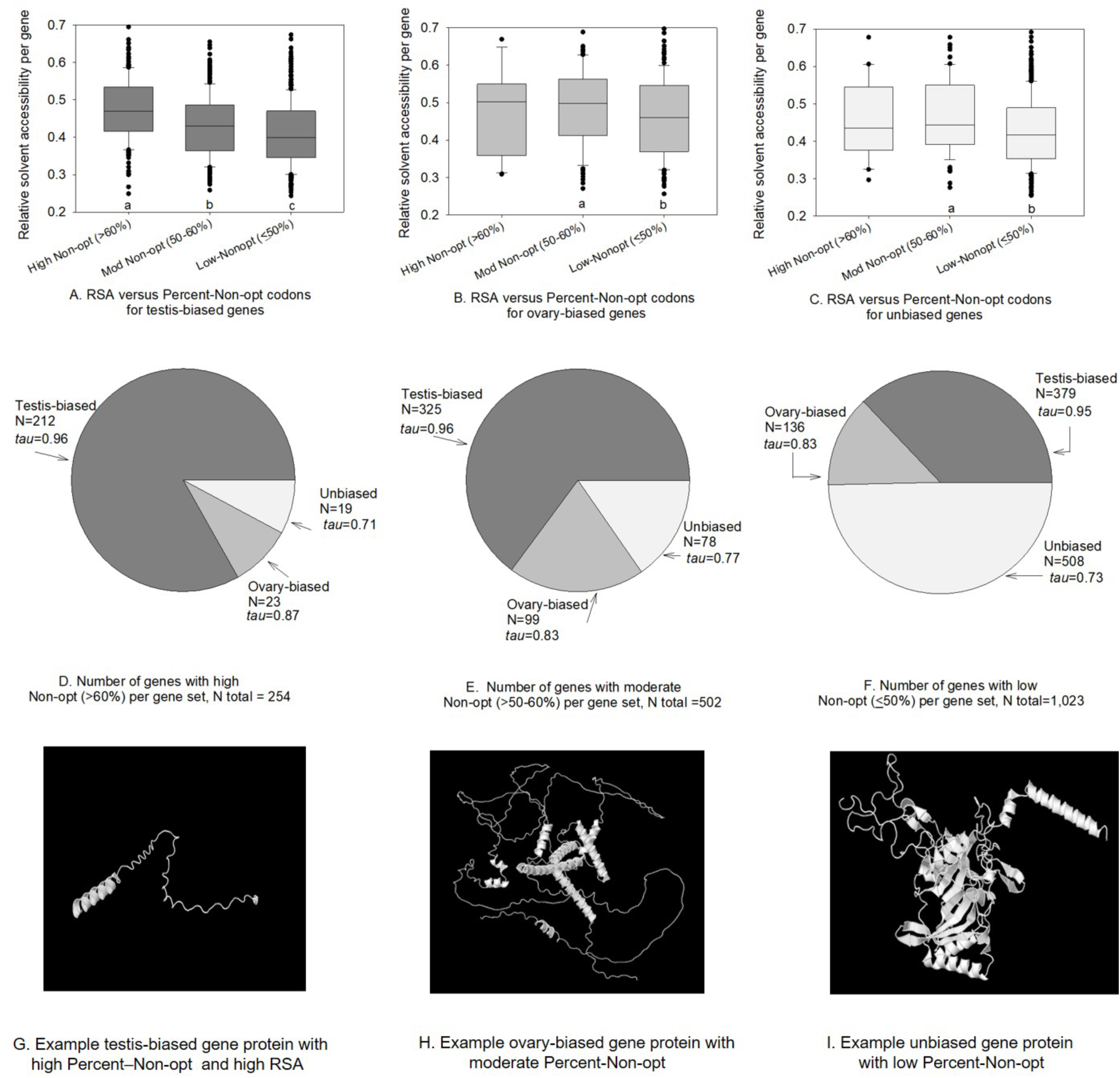
A-C: The relationships between high non-optimal codon use (>60% of amino acids), moderate non-optimal use (50 to ≤60%) and low optimal codon use (≤50%) and the relative solvent accessibility for: A) testis-biased genes; B) ovary-biased genes; and C) unbiased genes in *D. melanogaster*. Pie charts of the number genes with D) high non-optimal codon use; E) moderate non-optimal; and F) high optimal codon use, also showing the median values of *tau* across genes. Examples of AlphaFold2 predicted protein structures of a typical: G) testis-biased gene with high non-optimal codons and RSA; H) ovary-biased genes wth moderate non-optimal codons and RSA; and I) an unbiased gene with low non-optimal codons and RSA. In A-C, different letters below bars indicate a statistically signficant difference (MWU-test P<0.05)

Importantly, of the 255 genes (out of 1,779 highly expressed genes under study; Fig. S1A) within the highest Percent-Non-opt class (>60% non-optimal codons, Fig. 2D), 83% were testis-biased (N=212), while only 9.0% were ovary-biased (N=23) and 7.8% were unbiased (N=20; Fig. 2D). Thus, the highest degree of non-optimal codon use is most commonly observed in the testis-biased genes, and is much less common among ovary-biased and unbiased genes. Given that non-optimal codons, when they occur, tend to be found in higher RSA regions (Fig, 2A-C), this implies a concentration of non-optimal codons in the high RSA regions of testis-biased genes (i.e. non-optimal codons less common overall in ovary-biased or unbiased genes). Even when taking into account that there were more testis-biased genes than ovary-biased or unbiased genes, we find that 23.1% (212 of 916) of the testis-biased gene set belongs to the highest class bin of non-optimal codon use, versus only 8.9% (23 of 258) and 3.3% (20 of 605) of the ovary-biased and unbiased gene sets respectively. In terms of genes in the moderate Percent-Non-opt bin (Fig. 2E), testis-biased genes were most common in absolute numbers (N=325; that 35.4% of all testis-biased genes), with less than a third as many ovary-biased genes (N=99 of 258, 38.3% of the ovary genes), and even fewer unbiased genes (N=78; 12.9% of unbiased genes). In the lowest Percent-Non-opt bin (with the highest optimal codon use), unbiased genes were most strongly represented with 509 genes (84.1% of the unbiased gene set), while 389 were testis-biased (42.5% of testis-biased gene set), and 136 were ovary-biased genes (52.7% of the ovary-biased gene set; Fig. 2F). Overall, it is evident that a major portion of testis-biased genes were linked to the highest non-optimal codon use bin (Percent-Non-opt>60%), a group that also had very high protein-wide RSA (Fig. 2A-C). These RSA patterns add another layer of support (in addition to the results in Fig. 1) to the conclusion that the high non-optimal codon use in *D. melanogaster* genes is non-random, given that it is also negatively associated with the degree of protein folding in all gene sets under study (Fig. 2A-C), and that that the highest degree of non-optimal codon use (and thus high RSA) is especially common in testis-biased genes (Fig. 2A,D).

In Figs. 2G-I, we provide examples of the predicted structures of a testis-biased gene from the highest Percent-Non-opt bin, an ovary-biased gene from the moderate Percent-Non-opt gene bin, and an unbiased gene from the low Percent-Non-opt bin (note RSA variation within each bin, Fig. 2A-C). The testis-biased sperm/accessory gland-related gene protein Mst57Da (FBgn0011668), a relatively short protein, encodes 73.0% of its amino acids with non-optimal codons, and is predicted to be highly unstructured (Fig. 2G) with an average CDS-wide RSA of 0.72±0.09 (for an example of a similar weakly folded protein structure for a longer testis-biased gene, BG642163, in the highest Percent-Non-opt bin see Fig. S6). The ovary-biased gene protein Jabba (FBgn0259682), that is involved in early embryogenesis (Gramates et al. 2022), has 55.1% of its amino acids encoded by non-optimal codons, and shows more helical structures and a lower RSA value (0.50 ±0.01; Fig. 2H) than the testis-protein (Fig. 2G). The unbiased gene encoding ribosomal protein RpL3 (FBgn0020910) has 73.2% optimal codon use and an average RSA of 0.36 ±0.01 (Fig. 2I), and high levels of helical and beta-sheets illustrating its extensive degree of folding (NetsurfP predictions for all three proteins are in Fig. S7). In sum, these patterns exemplify the tendency for positive associations in non-optimal codon use and RSA.

Functional clustering shows that the 212 testis-biased genes in the highest Percent-Non-opt bin (>60% non-optimal codons, Fig. 2D) are enriched in genes involved in sperm-related functions (Table S6). Examples include genes encoding accessory gland proteins Acp54A1, Acp26Aa and Acp36DE, the testis-enriched gene *RpL10Aa,* the seminal fluid protein Sfp87B, and spermatogenesis genes *soti*, *Mst89B*, M*st57Da*, and other Mst genes (Table S7; gene functions from DAVID and FlyBase (Sherman et al. 2022; Gramates et al. 2022)). Sperm and accessory gland genes (Tables S6, S7; an example structure in Fig. 2G) are of particular interest to evolutionary biologists due to their key role in sexual reproductive success, and their often-rapid rates of protein sequence divergence, including in *Drosophila* (Torgerson et al. 2002; Civetta & Clark 2000; Dorus et al. 2006; Pitnick 1996). Overall, these testis-biased genes with the highest percentage of non-optimal codons were highly specialized to testicular functions, including accessory gland and sperm functions.

### Tissue specificity suggests a mechanism to accumulate non-optimal codons

Analyses of gene expression specificity across 59 tissues/stages in *D. melanogaster* (Table S2), revealed that testis-biased genes had exceptionally elevated *tau* values, with a median near the maximum possible value of 1 for genes in the high, moderate and low Percent-Non-opt bins (median values≥0.96, Fig. 2D-F; see Materials and Methods). Thus, the expression of these genes is highly specific to the testis. In turn, ovary-biased genes, had intermediate *tau* values (median between 0.83 to 0.87) among low, moderate and high Percent-Non-opt bins, while unbiased genes had the lowest *tau* (0.71 to 0.77) (Fig. 2D-F). Assuming expression breadth reflects a gene’s cross-tissue pleiotropy (Meisel 2011; Mank & Ellegren 2009; Assis et al. 2012), then testis-biased genes comprise the least pleiotropic gene set under study. The low pleiotropy of these testis-biased genes may allow them to be under less severe functional constraint than those involved in a wide range of tissues or pathways (Otto 2004; Meisel 2011; Dean & Mank 2016; Mank et al. 2008), possibly allowing them the flexibility to evolve the observed high non-optimal codon use (Figs. 1, 2). Moreover, sex-biased gene expression, including sex-specific expression (79.0% of testis-biased genes were sex-specific herein, Fig. S1 legend), may act as a means to limit sexual conflict (Ellegren & Parsch 2007; Wong & Holman 2023; Ingleby et al. 2014). Thus, we predict that non-optimal codon use in the testis-biased genes may typically be of little or no fitness consequence to the ovaries or other tissues, facilitating accumulation of non-optimal codons in testis genes to regulate gene expression for specialized testicular functions.

### All 18 Degenerate Amino Acids have Higher RSA Values When Encoded by the Primary Non-optimal Codon than by the Optimal Codon

Finally, we assessed the RSA value for each individual type of amino acid when it was encoded by the primary non-optimal codon (Table S5) versus when it was encoded by the optimal codon (Table S1) in the testis-biased genes, Our analyses of all amino acids encoded by testis-biased genes showed that for all 18 degenerate amino acids, the primary non-optimal codon was linked to a higher average RSA than the primary optimal codon (Fig. 3; sign test P<0.001; averages and standard errors are in Table S8). Thus, the elevated RSA of non-optimal codons included every degenerate amino acid, regardless of its maximum solvent accessibility and its polarity (which vary among amino acids (Tien et al. 2013)). When encoded by a non-optimal codon, residues were consistently located in less deeply folded protein regions, than when that same amino acid was encoded by its optimal codon. This result includes the primary non-optimal codons Thr-ACT, Ile-ATA, and Gly-GGA (Fig. 2B-D) and spans all two-fold, three-fold, four-fold and six-fold degenerate amino acids (Table S5). Thus, the favored use of the primary non-optimal codons versus the optimal codon in more disordered regions was universal across all 18 degenerate amino acids.

**Fig. 3.**
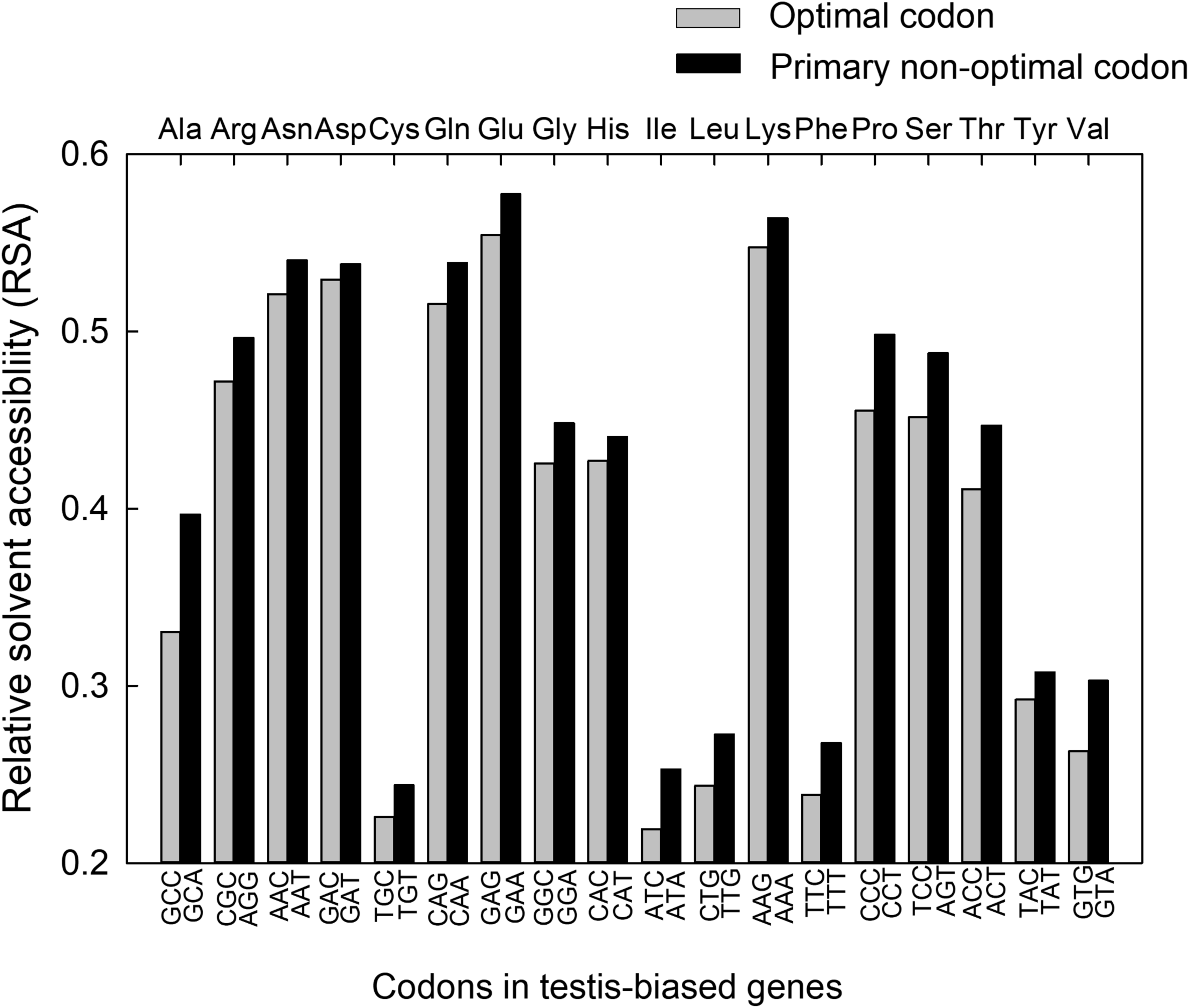
The average RSA of testis-biased genes when encoded by the primary non-optimal codon (defined in Table S5) versus the optimal codon (Table S1) for all 18 degenerate amino acids. Results are from all amino acids and codons in testis-biased genes (N=299,817 amino acids). In total, 18 of 18 degenerate amino acids had higher average RSA when encoded by the non-optimal codon; a sign test of the non-optimal versus optimal codon across all 18 amino acids had P<0.001. The standard errors were very small (all <0.005) and are provided in Table S8.

In addition, the primary non-optimal codons, which all exhibited elevated RSA compared to optimal codons, included those with few and those with plentiful matching tRNA genes (i.e. with Non-opt_low-tRNAs_ and Non-opt_high-tRNAs_ statuses (Fig. 3, Table S1, Table S5; optimal codons included both Opt_high-tRNAs_ and Opt_wobble_ statuses). The rarity of exact matching tRNAs for Non-opt_low-tRNAs_ codons (Table S1, Table S5, Fig. 3) may make these types of codons poised to slow translation, and allow proper co-translational protein folding (Brule & Grayhack 2017; Stein & Frydman 2019), including in disordered regions (Zhou et al. 2015). However, the high ratio of tRNA genes to codon use for Non-opt_high-tRNAs_ codons suggests that this class of codons likely has different functions, possibly in the rapid translation of mRNA segments containing the codons, or the up-translation of any mRNAs with high use of these (otherwise) rare codons within a cellular mRNA pool (Whittle et al. 2021; Torrent et al. 2018; Whittle et al. 2019). Thus, despite potentially having distinct functions in slowing translation (Non-opt_low-tRNAs_) or in targeted enhanced translation of mRNAs (Non-opt_high-tRNAs)_) (Brackley et al. 2011; Zhou et al. 2015; Stein & Frydman 2019; Brule & Grayhack 2017; Whittle et al. 2021; Liu et al. 2021; Fu et al. 2016), the two types of non-optimal codons (Table S1), are each consistently favored over optimal codons in less-structured protein segments (Fig. 3). Based on these findings, we speculate that the combined use of primary Non-opt_low-tRNAs_ and Non-opt_high-tRNAs_ codons (Fig. 3) may respectively act to slow down and speed up local rates of translation within specific types of mRNAs (for instance, in highly transcribed mRNAs that have ameliorated use these otherwise rare codons, such as testis and sperm genes (Table S5, Fig. 3, and/or in some regions of ovary-biased and unbiased genes (Fig. 2B, C), thereby providing a means to control translational pacing and potentially allowing proper protein conformation and functionality.

### The Potential Role of Selection in Shaping Non-optimal Codon Use

Some of the non-optimal codon use patterns observed here may reflect a history of relaxation of constraint, particularly among the non-primary non-optimal codons per amino acid (Fig. 1, Table S5). However, several lines of evidence support a hypothesis that selection has at least partly contributed to the use of a subset of preferred (primary) non-optimal codons. First, we *a priori* identified optimal and non-optimal codons from analysis of the most highly expressed genes in the *D. melanogaster* genome (Table S1), which are most apt to experience translation-related selection pressures (including tRNA abundance pressures), using an approach parallel to that historically used to identify optimal codons (Duret & Mouchiroud 1999; Whittle & Extavour 2016; Whittle et al. 2021, 2007; Duret 2000; Akashi 2001; Wang et al. 2011; Whittle et al. 2019). This *a priori* analysis automatically (in Table S1) controls for codon use in lowly and moderately expressed genes, wherein selection is weak or absent and neutral processes tend to be prevalent (Wang et al. 2011; Whittle et al. 2019, 2021). Second, a specific subset of non-optimal codons, typically one specific codon per amino acid, exhibited much higher use than the other sister non-optimal codons in highly transcribed testis-biased genes (Fig. 1, Table S5). We posit that this pattern, remarkably observed for 18 of 18 degenerate amino acids (Table S5), is unlikely to be explained by a fully neutral process, whereby all non-optimal codons would be expected to have similar, or arbitrary use patterns. Third, the same primary non-optimal codon was identified when using the ovary-biased genes and when using the unbiased genes as the reference dataset to compare with testis-biased genes (Fig. 1, Table S5); the ovary-biased and unbiased groups are mutually exclusive gene sets, but the same primary non-optimal codon per amino acid was nevertheless identified in in the testis-biased genes. Fourth, the primary (preferred) non-optimal codon for amino acids ended in different nucleotides (A, T, G; Fig. 1, Table S5) and thus a mutation-bias or gene conversion process (e.g., favoring G) associated with high gene expression in the testis is a poor explanation for this pattern. Fifth, our findings that 18 of 18 degenerate amino acids in *D. melanogaster* showed preferential location of the primary non-optimal codon in more disordered regions over its sister optimal codon (higher RSA, Fig. 3), infers a remarkably consistent pattern of non-arbitrary codon use among sister codons, across every degenerate amino acid, again unlikely to be the result of a neutral process. Further, we showed this pattern was observed for non-optimal codons that we *a priori* identified that had enhanced use in highly expressed genes (Table S1), which are most apt to experience translation-related selection pressures (as compared to low or moderately expressed genes (similar to optimal codons in highly expressed genes (Duret & Mouchiroud 1999; Whittle & Extavour 2016; Whittle et al. 2021, 2007; Duret 2000; Akashi 2001; Wang et al. 2011; Whittle et al. 2019)). While neutral processes are apt to play a role in shaping the use of the non-primary non-optimal codons per amino acid for highly expressed testis-biased and ovary-biased genes (for those amino acids with three or more codons and thus, more than one non-optimal codon, Fig. 1), we propose the collective evidence best supports a model whereby one (or rarely two) non-optimal codon is favored over others, and that use of these codons tends to be preferentially located in genes and gene regions that encode disordered proteins (Fig. 2, Fig. 3).

### Conclusions and Future Directions

The collective evidence herein supports the hypothesis that selection has contributed to shaping the use of a specific subset of (preferred) non-optimal codons. Non-optimal codons are non-arbitrarily used in sex-related genes in *D. melanogaster*, and certain non-optimal codons are favored over others. This pattern is particularly prevalent in testis-biased genes, and suggests that certain (primary, or preferred) non-optimal codons could have functions in regulating translation.

Future studies should assess the prevalence and evidence for selection for non-optimal codon use in other multicellular organisms, including a system whereby non-optimal codons are more common in the ovaries (Whittle & Extavour 2017) or in other specific organ systems. In addition, investigation of the placement of all four categories of codons studied herein, namely Opt_high-tRNAs_, Opt_wobble,_ Non-opt_low-tRNAs_ and Non-opt_high-tRNAs_, with respect to the starting ramp of translation, versus middle and terminal CDS regions, and especially with respect to the RSA in disordered regions, will help reveal whether these codons may cause localized pacing (fast/slow) of translation (Quax et al. 2015; Stein & Frydman 2019; Miller et al. 2019) in *D. melanogaster* and other multicellular models. Experimental studies to quantify translation rates and protein folding of testis genes with extreme use of Non-opt_high-tRNAs_ codons (Fig. S4, Table S5; cf. (Zhou et al. 2015; Allen et al. 2022)) may further reveal their putative roles in the rapid translation of specific mRNAs in a cellular mRNA pool or of mRNA regions (Torrent et al. 2018; Whittle et al. 2021), while investigations of those genes with extreme use of Non-opt_low-tRNAs_ codons (Fig. S4) may inform on their function in the slowed translation and precise protein conformation in testis-involved genes, as inferred for circadian genes (Fu et al. 2016; Zhou et al. 2015; Liu et al. 2021). Additional data on the tRNA populations in the testes and ovaries in this species as the technology improves (Guimaraes et al. 2020) (discussed in Supplementary Text File S1), will also help determine whether or how there is variation in tRNA populations among the gonads, beyond that predicted by tRNA gene frequency (Fig. 2). Future research should also evaluate whether younger testis-biased genes have greater flexibility to evolve a set of preferred non-optimal codons with functional roles (*cf.*(Allen et al. 2022)), and whether non-optimal codon use may have particular roles in shorter genes (as testis-biased genes tended to be short herein, Fig. S3, Fig. 2). Finally, *in vitro* or *in vivo* studies that replace non-optimal codons with optimal codons (Zhou et al. 2015; Fu et al. 2016; Allen et al. 2022), and vice-versa, to evaluate their effects on protein structure and functions for testis-biased and ovary-biased genes studied here, may provide further insights into the potential specific protein biophysical roles of the non-optimal codons in this species. As part of such analyses, an evaluation of altered Non-opt_low-tRNAs_ and Non-opt_high-tRNAs_ statuses within disordered protein regions may reveal their individual roles on the pacing of translation, protein conformation and protein functionality.

## Materials and Methods

### Genomic Data and Expression Analyses

We downloaded the genome of *D. melanogaster* (13,986 protein coding genes) version 6.50 (Gramates et al. 2022). For our analysis, we extracted the longest CDS per gene (Whittle et al. 2021, 2019; Sandmann et al. 2011). To identify sex-biased genes, we used the modEncode database that contains expression data (Graveley et al. 2011; Gramates et al. 2022) for a wide range of *D. melanogaster* tissue types and developmental stages (N=59 tissues/stages studied herein, Table S2 (Li et al. 2014; Gramates et al. 2022; Graveley et al. 2011)), and that we also used to measure *tau* (Yanai et al. 2005) across all tissues. We identified highly expressed sex-biased genes as those with ≥100RPKM and ≥5-fold sex-bias in one sex versus the other in the contrast of 4-day mated testes versus 4-day mated ovaries, and unbiased genes as those with <5-fold bias (and ≥100RPKM in at least one gonad). For additional rigor, we also identified sex-biased genes using male accessory glands versus virgin ovaries (Table S2). Further details on identification of sex-biased genes and calculations of expression specificity using *tau* ((Yanai et al. 2005)(described previously in (Whittle & Extavour 2023)) using the data in Table S2, are provided in Supplementary Text File S1.

### Codon Use and **Δ**RSCU

The organism-wide RSCU values (Table S1) for *D. melanogaster* were obtained from the Codon Statistics Database (CSD) (Subramanian et al. 2022). From these, we calculated the ΔRSCU_Ribosome-All_ =RSCU_Ribosomal genes_ -RSCU_All genes_ (using the concatenated genes per dataset), which is a more conservative version of the approach using the highest and least expressed genes (Wang et al. 2011; Whittle et al. 2007; Cutter et al. 2006; Ingvarsson 2008; Duret & Mouchiroud 1999). We identified the largest positive ΔRSCU_Ribosome-All_ values as the optimal codon (and each was also identified as optimal at the CSD, with P-values<0.05). All other codons per amino acid were classed as non-optimal, given their large disparity in use from the optimal codon (see Supplementary Text File S1 section “Genome Datasets and Gonadal Expression Analyses”). The tRNA gene counts in the *D. melanogaster* genome were determined using the GtRNA database (http://gtrnadb.ucsc.edu) (Chan & Lowe 2016) which utilizes tRNAscan-SE to extract tRNAs from the genome (Lowe & Chan 2016; Chan & Lowe 2019). CAI values that reflect similarity of codon use to the highly expressed ribosomal gene set, and the Fop values, were obtained from the CSD (Subramanian et al. 2022).

For the study of differences codon use among sex-biased genes, we determined herein the RSCU values and the codon raw counts per codon for every testis-biased, ovary-biased and unbiased gene using Codon W (Peden 1999). For each of the studied 59 codons encoding a degenerate amino acid, the RSCU values were used to calculate ΔRSCU_Testis-Ovary_=average RSCU_Testis-biased genes_ – average RSCU_Ovary-biased genes_ and the ΔRSCU_Testis-Unbiased_=average RSCU_Testis-biased genes_ – average RSCU_Unbiased genes_. The RSCU values per codon were compared across all genes per contrast using t-tests to obtain P-values (Whittle et al. 2007). The codon counts per gene under study (for each testis-biased, ovary-biased and unbiased gene) were used to calculate the Percent-Non-opt codon use per gene (relative to all optimal and non-optimal codons per gene), based on the non-optimal and optimal codon identities that were defined in Table S1 (see also Supplementary Text File S1). The RSCU values used to calculate ΔRSCU for sex-biased and unbiased genes are shown in Tables S9 and S10.

### Protein Folding, Relative Solvent Accessibility and Codon Use in the Sex-biased Genes

To determine the RSA per amino acid we used NetsurfP v3.0 set to default parameters (Hoie et al. 2022). This program is a machine-learning based tool proven effective for determining RSA and other structural data of proteins (Hoie et al. 2022; Klausen et al. 2019). NetsurfP v3.0 has been widely used to characterize protein folding and structures (Hoie et al. 2022; Klausen et al. 2019), and its RSA calculations have been shown to be equivalent to those predicted by other top predictor tools such as SPOT-1D-Single (Hoie et al. 2022). We also found strong correlations using protein structures from the Protein Data Base and RSA from GETAREA (Braun 1998) (data not shown). We determined the type of secondary structure that each amino acid was located within, namely a helix, string (β-string/sheet) or a random coil, and the disorder scores per amino acid, using NetsurfP 3.0 (Hoie et al. 2022).

Protein folding predictions for the visualization of structures in Fig. 2G-I were determined using Alphafold2 (Juniper et al. 2021), available for the *D. melanogaster* proteins (Uniprot proteome Identity UP000000803, taxon identity= 7227) from the AlphaFold2 database (https://alphafold.ebi.ac.uk) (Varadi et al. 2022). Visualization was conducted using Geneious Prime (Kearse et al. 2012).

For Fig. 3, using all concatenated testis-biased genes, the RSA was determined for each type of amino acid (18 degenerate amino acids) when it was encoded by the preferred non-optimal codon and when the same amino acid was encoded by the optimal codon, across all amino acid sites. Then, the average RSA was determined for all cases where the amino acid was encoded by the preferred non-optimal codon and when encoded by the optimal codon (averages and standard errors are provided in Table S8).

### GO Functions

Gene ontology and functional analysis was performed using DAVID (Sherman et al. 2022).

## Data Availability

All data used in this study are publicly available. The locations of the data are provided in the Materials and Methods.

## Acknowledgements

The authors thank members of the Extavour lab for valuable discussions. C. Extavour is an Investigator of the Howard Hughes Medical Institute, which supported this work. The valuable comments from two anonymous Reviewers are appreciated.

