## Supplementary Information for "Evidence for non-optimal codon choice in highly transcribed sex-biased genes of *Drosophila melanogaster*"

**Contents**

Figs. S1 to S7

Tables S1 to S10

Supplementary text (Results and Discussion; Materials and Methods)

References for SI citations

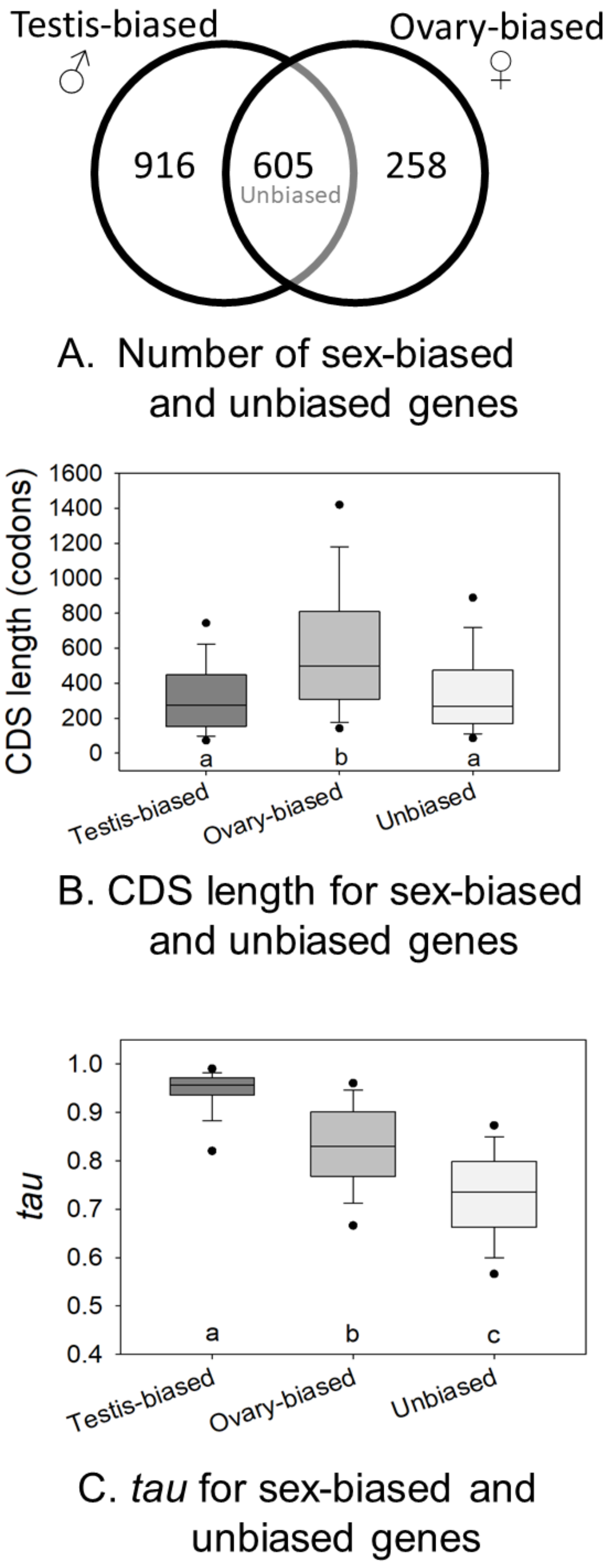

**Fig. S1.** (A) The number of sex-biased and unbiased genes under study. (B) CDS lengths per gene set. (C) the *tau* values per gene set. For A, the numbers of genes with testis-specific and ovary-specific expression (0 RPKM) with respect to the opposite sexual tissue was 724 (79.0%) and 19 (7.4%) respectively. Different letters under bars in B and C indicate a statistically significant difference with MWU-test $P<0.05$.

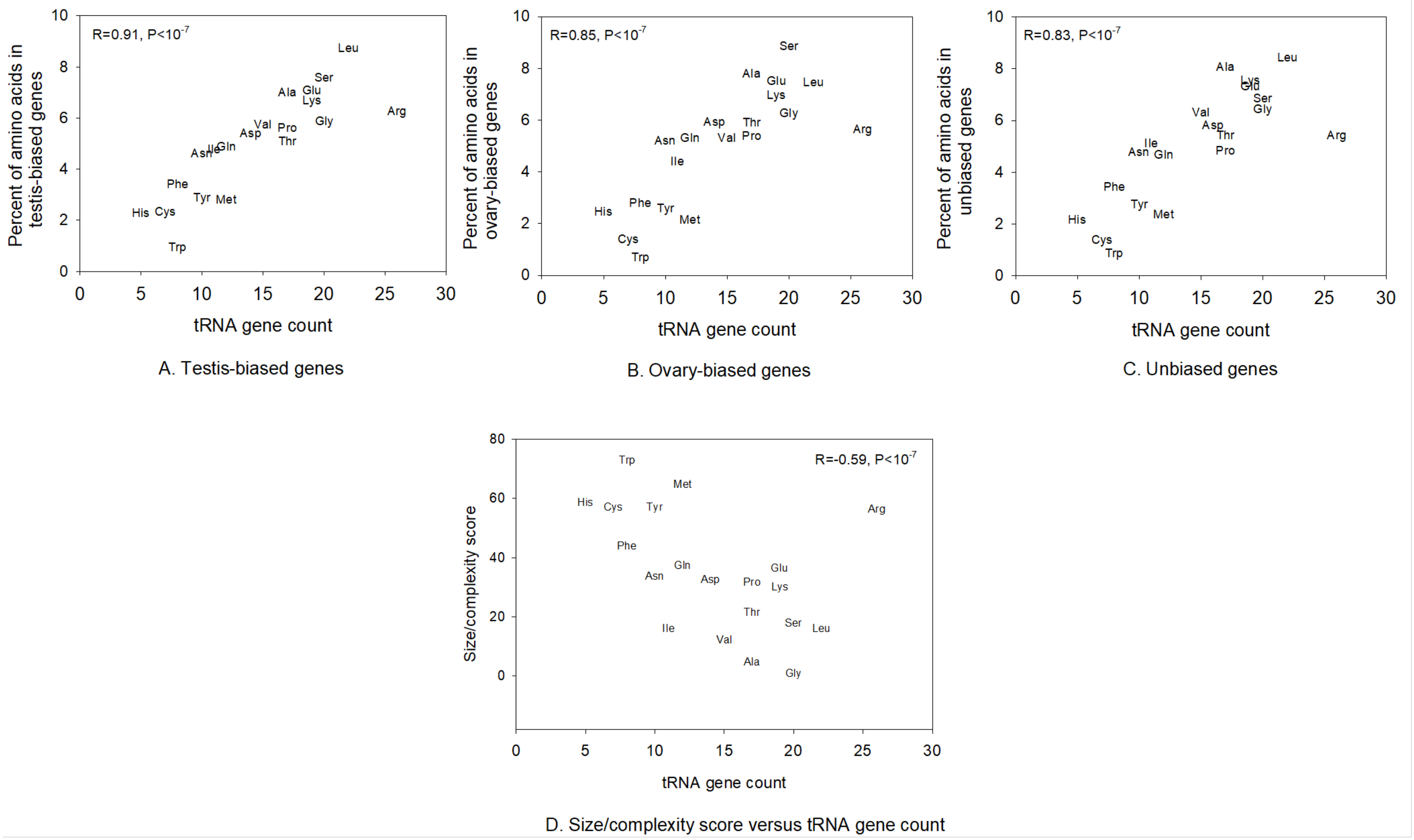

**Fig. S2.** The relationship between the frequency of amino acid use (percent of all concatenated amino acids per gene set) and tRNA genes in the genome of *D. melanogaster*. (A) testis-biased genes. (B) ovary-biased genes. (C) unbiased genes. (D) The association between the size/complexity score per amino acid and tRNA gene counts in the genome. The Spearman Rank correlations are shown. The number of amino acids for testis-biased genes was N=299,817, for ovary-biased genes N=155,282 amino acids, and for unbiased genes was N=218,580 amino acids.

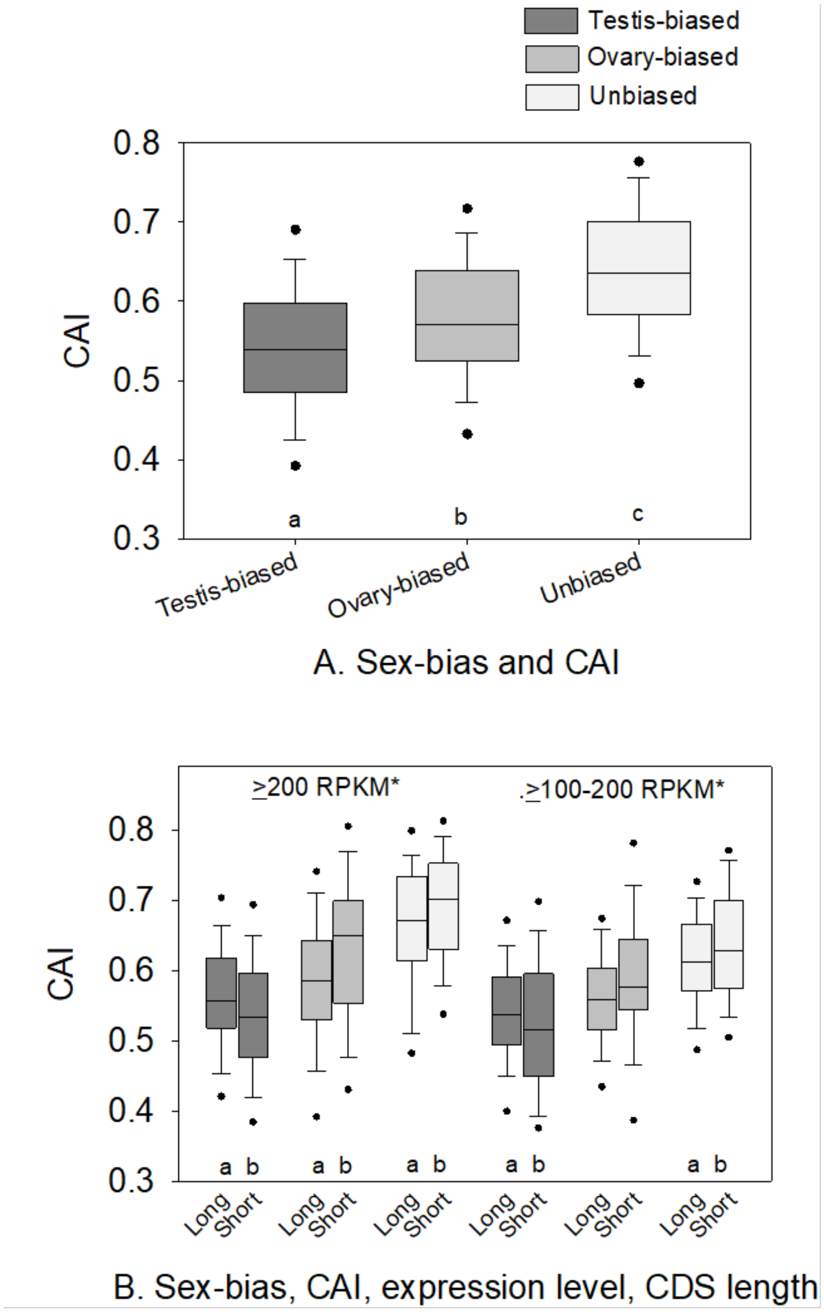

**Fig. S3.** Box plots of CAI for *D. melanogaster* sex-biased gene sets under study. (A) CAI of testis-biased, ovary-biased and unbiased genes. (B) CAI for testis-biased, ovary-biased and unbiased genes subdivided by gene expression level, and by CDS length. In A, different letters among bars indicates statistical significance using ranked ANOVA and Dunn's paired contrast ($P<0.05$). In panel B, different letters under each set of paired bars indicates CAI between short and long CDS had MWU-test $P<0.05$. *Indicates a difference in CAI between the high and low expressed genes for the testis-biased genes, the ovary-biased genes, and for the unbiased genes (MWU-test $P<0.05$; grouped genes across both length classes).

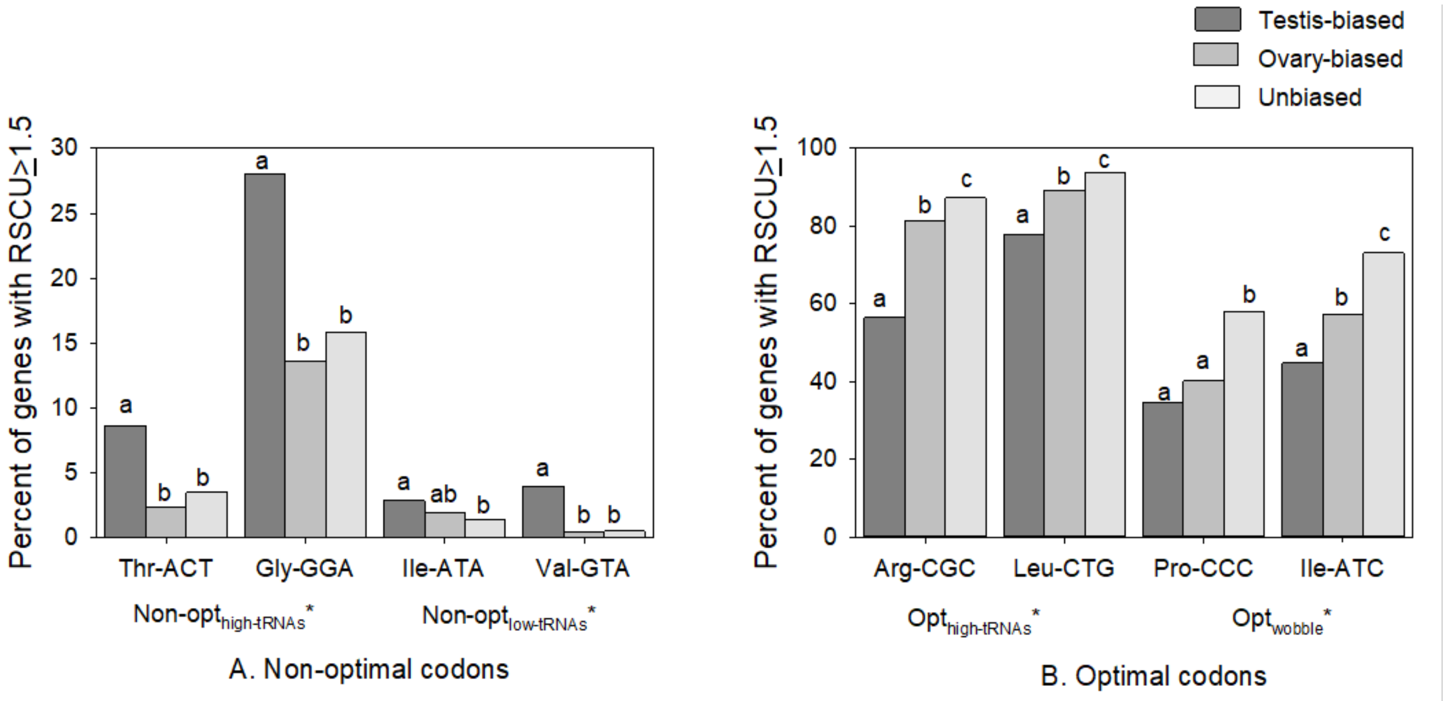

**Fig. S4.** Examples of non-optimal and optimal codons with very high use (RSCU≥1.5) in sex-biased and unbiased genes. (A) Four non-optimal codons, including two with high and two with a low number of matching tRNA genes. (B) optimal codons, including two with high numbers of matching tRNA genes and two requiring wobble tRNAs. In (A) and (B), different letters among the three bars for each codon indicates a statistically significant difference using Chi-square tests (P<0.05).* (asterisk) in (A) indicates percentages were higher for each Non-opt$_{high\text{-}tRNAs}$ codon versus each comparable Non-opt$_{lowtRNAs}$ codon for paired contrasts between testis-biased, ovary-biased and between unbiased genes (Chi-square P<0.05, with the only exception being between Thr-ACT and Ile-ATA for the ovary-biased genes P>0.05). * (asterisk) in (B) indicates all percentages were higher for each Opt$_{high\text{-}tRNAs}$ codon than each Opt$_{wobble}$ codon in paired contrasts between testis-biased genes, between ovary-biased genes and between unbiased genes (Chi-square P<0.05). The N for testis-biased genes =916, ovary-biased =258 and unbiased=605 genes. Note the Y-axis scales are different in (A) and (B).

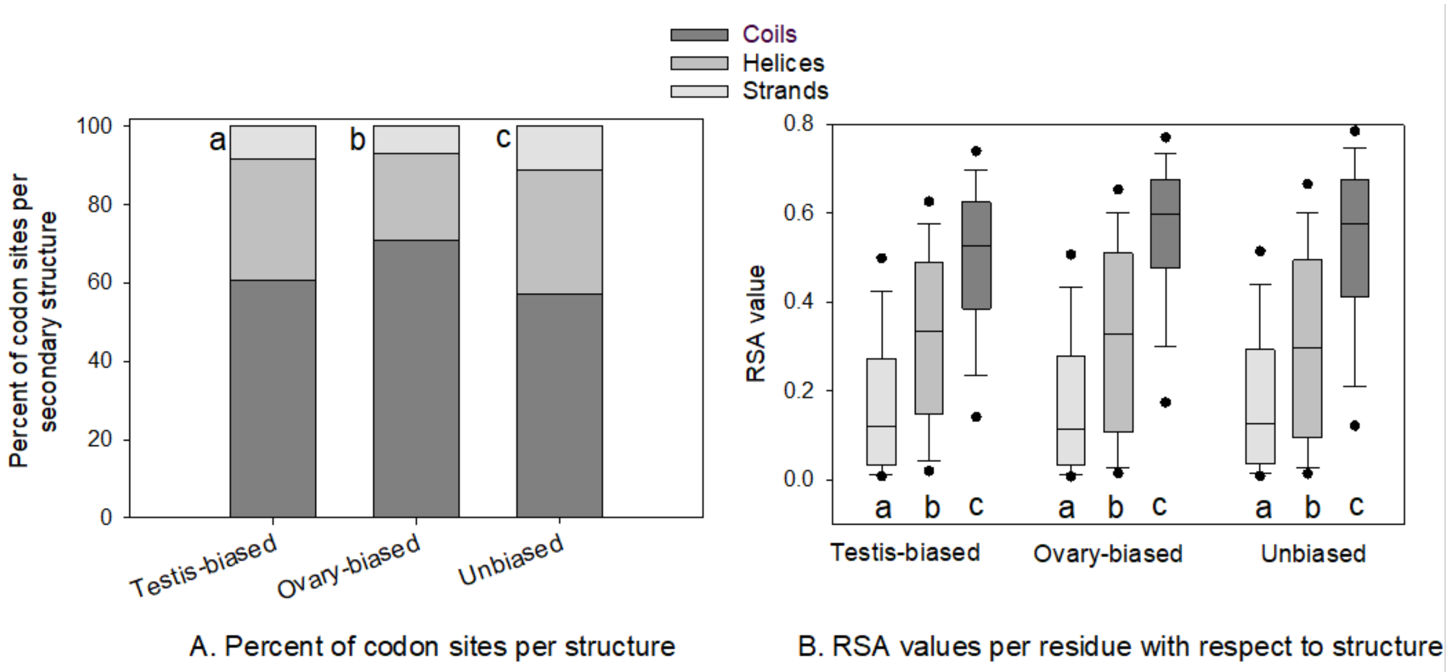

**Fig. S5.** (A) The percent of amino acids in sex-biased and unbiased genes located in sheets (strands), helices, and coils. (B) The RSA values for sex-biased and unbiased genes for all amino acids across all proteins. In (A), the percent in stands, helices and coils differed within all three gene sets (Chi-square tests $P<0.05$). In addition, a higher percemt of codons were located in coils for ovary-biased genes than testis-biased and unbiased genes (Chi-square test $P<0.05$). In (B), different letters within testis-biased, ovary-biased and unbiased genes indicates a statistically signficant difference in RSA using MWU-tests ($P<0.05$). All sites were concatenated across genes for each sex-biased gene category.

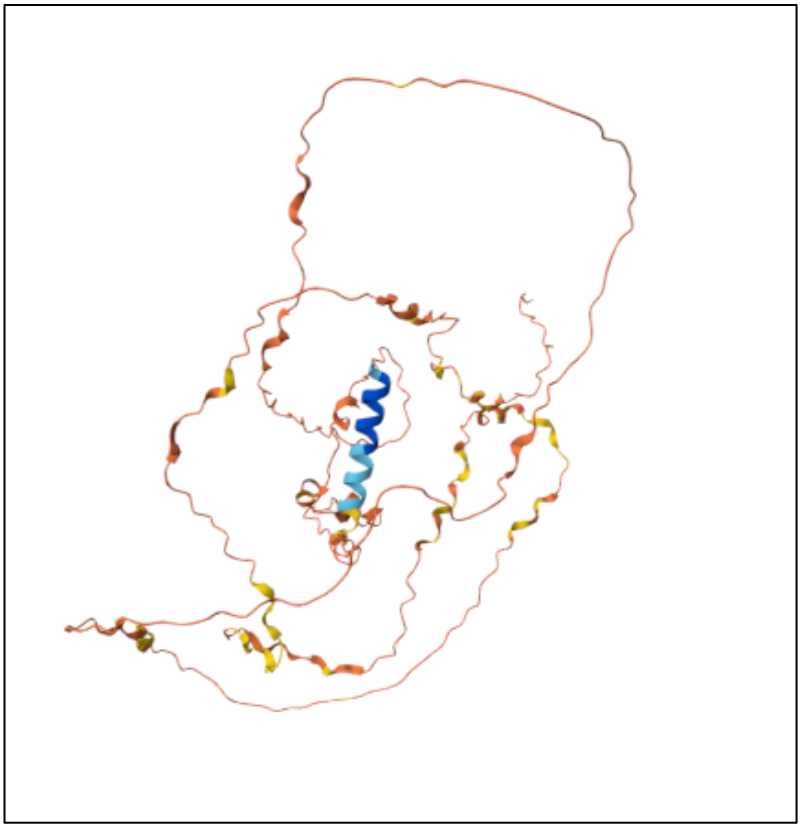

**Fig. S6.** An example of the predicted structure of a relatively long testis-bised gene protein (415 amino acids, protein ID=BG642163, Flybase ID= FBgn0083938) in the high Percent-Non-opt bin. Stucture is from Alpha-Fold (Jumper et al. 2021; Fleming et al. 2025) .

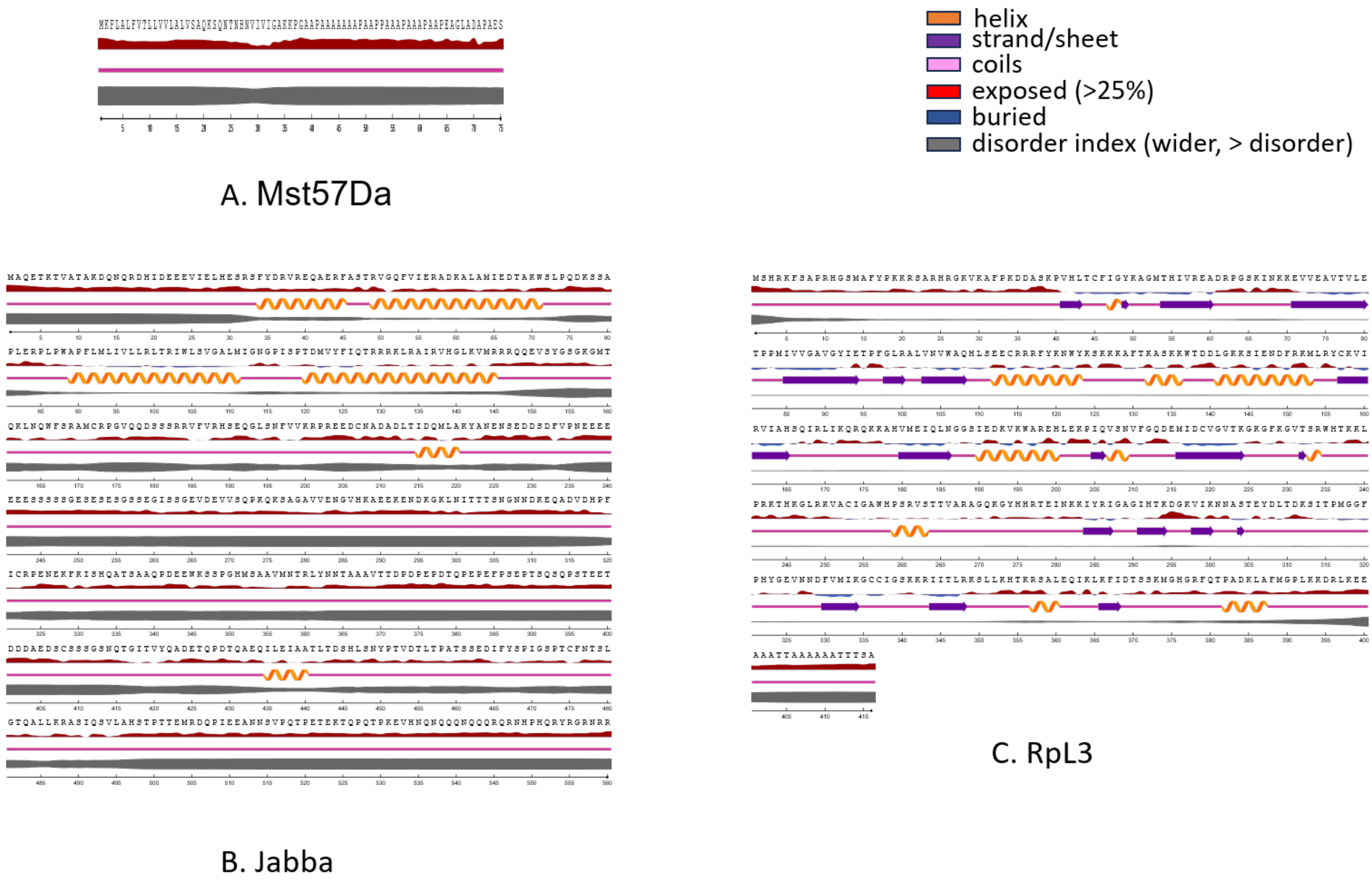

**Fig. S7.** Examples of the output of NetsurfP 3.0 (Hoie et al. 2022) for three genes provided in Fig. 3 G-I. RSA was determined for each amino acid based on its surface exposure.

**Table S1.** The organismal optimal and non-optimal codons for *D. melanogaster*. The ΔRSCU was determined between ribosomal genes versus all genes in the genome (ΔRSCU$_{\text{Ribosome-All}}$) and the number of exact matching tRNA genes were obtained from the GtRNA database (Chan and Lowe 2016). The codon statuses are defined in the main text and are as follows: Opt$_{\text{high-tRNAs}}$ = an optimal codon with abundant tRNA genes (≥5). Opt$_{\text{wobble}}$=an optimal codon requiring a wobble tRNA for translation, Non-opt$_{\text{low-tRNAs}}$= a non-optimal codon with few or no matching tRNA genes (<5), Non-opt$_{\text{high-tRNAs}}$= a non-optimal codon with a high number of matching tRNA genes (≥5). Optimal codons are in bold.

| **Amino Acid** | **Codon** | **Dmel ribosomal RSCU** | **All genes RSCU** | **ΔRSCU Dmel** | **Optimal and non-optimal status** | **No. of tRNAs** | |
|---|---|---|---|---|---|---|---|
| | | | | | | Anti-codon | No. tRNAs |
| Ala | GCT | 0.85 | 0.79 | +0.06 | Non-opt$_{\text{high-tRNAs}}$ | AGC | 12 |
| **Ala** | **GCC** | **2.14** | **1.75** | **+0.39** | **Opt$_{\text{wobble}}$** | **GGC** | **0** |
| Ala | GCA | 0.44 | 0.71 | -0.27 | Non-opt$_{\text{low-tRNAs}}$ | TGC | 2 |
| Ala | GCG | 0.58 | 0.75 | -0.17 | Non-opt$_{\text{low-tRNAs}}$ | CGC | 3 |
| Arg | CGT | 1.42 | 0.93 | +0.49 | Non-opt$_{\text{high-tRNAs}}$ | ACG | 10 |
| **Arg** | **CGC** | **2.67** | **1.86** | **+0.81** | **Opt$_{\text{wobble}}$** | **GCG** | **0** |
| Arg | CGA | 0.48 | 0.95 | -0.47 | Non-opt$_{\text{high-tRNAs}}$ | TCG | 10 |
| Arg | CGG | 0.56 | 0.91 | -0.35 | Non-opt$_{\text{low-tRNAs}}$ | CCG | 0 |
| Arg | AGA | 0.31 | 0.61 | -0.3 | Non-opt$_{\text{low-tRNAs}}$ | TCT | 3 |
| Arg | AGG | 0.57 | 0.74 | -0.17 | Non-opt$_{\text{low-tRNAs}}$ | CCT | 3 |
| Asn | AAT | 0.63 | 0.93 | -0.30 | Non-opt$_{\text{low-tRNAs}}$ | ATT | 0 |
| **Asn** | **AAC** | **1.37** | **1.07** | **+0.30** | **Opt$_{\text{high-tRNAs}}$** | GTT | **10** |
| Asp | GAT | 0.94 | 1.08 | -0.14 | Non-opt$_{\text{low-tRNAs}}$ | ATC | 0 |
| **Asp** | **GAC** | **1.06** | **0.92** | **+0.14** | **Opt$_{\text{high-tRNAs}}$** | **GTC** | **14** |
| Cys | TGT | 0.39 | 0.61 | -0.22 | Non-opt$_{\text{low-tRNAs}}$ | ACA | 0 |
| **Cys** | **TGC** | **1.61** | **1.39** | **+0.22** | **Opt$_{\text{high-tRNAs}}$** | GCA | **7** |
| Gln | CAA | 0.47 | 0.63 | -0.16 | Non-opt$_{\text{low-tRNAs}}$ | TTG | 4 |
| Gln | **CAG** | **1.53** | **1.37** | **+0.16** | **Opt$_{\text{high-tRNAs}}$** | CTG | **8** |
| Glu | GAA | 0.54 | 0.7 | -0.16 | Non-opt$_{\text{high-tRNAs}}$ | TTC | 6 |
| Glu | **GAG** | **1.46** | **1.3** | **+0.16** | **Opt$_{\text{high-tRNAs}}$** | **CTC** | **13** |
| Gly | GGT | 0.99 | 0.86 | +0.13 | Non-opt$_{\text{low-tRNAs}}$ | ACC | 0 |

| | | | | | | | |
|---|---|---|---|---|---|---|---|
| Gly | **GGC** | **1.81** | **1.66** | **+0.15** | **Opt$_{high-tRNAs}$** | **GCC** | **14** |
| Gly | GGA | 1.02 | 1.18 | -0.16 | Non-opt$_{high-tRNAs}$ | TCC | 6 |
| Gly | GGG | 0.17 | 0.31 | -0.14 | Non-opt$_{low-tRNAs}$ | CCC | 0 |
| His | CAT | 0.65 | 0.81 | -0.16 | Non-opt$_{low-tRNAs}$ | ATG | 0 |
| His | **CAC** | **1.35** | **1.19** | **+0.16** | **Opt$_{high-tRNAs}$** | **GTG** | **5** |
| Ile | ATT | 0.92 | 1.04 | -0.12 | Non-opt$_{high-tRNAs}$ | AAT | 9 |
| Ile | **ATC** | **1.77** | **1.35** | **+0.42** | **Opt$_{wobble}$** | **GAT** | **0** |
| Ile | ATA | 0.31 | 0.62 | -0.31 | Non-opt$_{low-tRNAs}$ | TAT | 2 |
| Leu | TTA | 0.2 | 0.32 | -0.12 | Non-opt$_{low-tRNAs}$ | TAA | 4 |
| Leu | TTG | 0.91 | 1.1 | -0.19 | Non-opt$_{low-tRNAs}$ | CAA | 4 |
| Leu | CTT | 0.47 | 0.62 | -0.15 | Non-opt$_{low-tRNAs}$ | AAG | 4 |
| Leu | CTC | 0.9 | 0.91 | -0.01 | Non-opt$_{low-tRNAs}$ | GAG | 0 |
| Leu | CTA | 0.36 | 0.57 | -0.21 | Non-opt$_{low-tRNAs}$ | TAG | 2 |
| Leu | **CTG** | **3.15** | **2.48** | **+0.67** | **Opt$_{high-tRNAs}$** | **CAG** | **8** |
| Lys | AAA | 0.37 | 0.63 | -0.26 | Non-opt$_{high-tRNAs}$ | TTT | 6 |
| **Lys** | **AAG** | **1.63** | **1.37** | **+0.26** | **Opt$_{high-tRNAs}$** | **CTT** | **13** |
| Met | ATG | 1 | 1 | 0 | | CAT | 12 |
| Phe | TTT | 0.52 | 0.79 | -0.27 | Non-opt$_{low-tRNAs}$ | AAA | 0 |
| **Phe** | **TTC** | **1.48** | **1.21** | **+0.27** | **Opt$_{high-tRNAs}$** | **GAA** | **8** |
| Pro | CCT | 0.48 | 0.54 | -0.06 | Non-opt$_{high-tRNAs}$ | AGG | 7 |
| **Pro** | **CCC** | **1.82** | **1.29** | **+0.53** | **Opt$_{wobble}$** | **GGG** | **0** |
| Pro | CCA | 0.7 | 1.03 | -0.33 | Non-opt$_{high-tRNAs}$ | TGG | 5 |
| Pro | CCG | 1 | 1.14 | -0.14 | Non-opt$_{high-tRNAs}$ | CGG | 5 |
| Ser | TCT | 0.53 | 0.53 | 0 | Non-opt$_{high-tRNAs}$ | AGA | 8 |
| Ser | **TCC** | **2.02** | **1.4** | **+0.62** | **Opt$_{wobble}$** | **GGA** | **0** |
| Ser | TCA | 0.41 | 0.59 | -0.18 | Non-opt$_{low-tRNAs}$ | TGA | 2 |
| Ser | TCG | 1.17 | 1.16 | 0.01 | Non-opt$_{low-tRNAs}$ | CGA | 4 |
| Ser | AGT | 0.5 | 0.87 | -0.37 | Non-opt$_{low-tRNAs}$ | ACT | 0 |
| Ser | AGC | 1.37 | 1.45 | -0.08 | Non-opt$_{high-tRNAs}$ | GCT | 6 |
| Thr | ACT | 0.57 | 0.71 | -0.14 | Non-opt$_{high-tRNAs}$ | AGT | 8 |
| **Thr** | **ACC** | **2.03** | **1.47** | **+0.56** | **Opt$_{wobble}$** | **GGT** | **0** |

| | | | | | | | |
|---|---|---|---|---|---|---|---|
| Thr | ACA | 0.52 | 0.82 | -0.3 | Non-opt$_{high\text{-}tRNAs}$ | TGT | 6 |
| Thr | ACG | 0.88 | 1 | -0.12 | Non-opt$_{low\text{-}tRNAs}$ | CGT | 3 |
| Trp | TGG | 1 | 1 | 0 | | CCA | 8 |
| Tyr | TAT | 0.56 | 0.77 | -0.21 | Non-opt$_{low\text{-}tRNAs}$ | ATA | 0 |
| **Tyr** | **TAC** | **1.44** | **1.23** | **+0.21** | **Opt$_{high\text{-}tRNAs}$** | **GTA** | **10** |
| Val | GTT | 0.65 | 0.77 | -0.12 | Non-opt$_{high\text{-}tRNAs}$ | AAC | 6 |
| Val | GTC | 1.25 | 0.91 | +0.34 | Non-opt$_{low\text{-}tRNAs}$ | GAC | 0 |
| Val | GTA | 0.27 | 0.45 | -0.18 | Non-opt$_{low\text{-}tRNAs}$ | TAC | 2 |
| **Val** | **GTG** | **1.84** | **1.86** | **-0.02** | **Opt$_{high\text{-}tRNAs}$** | **CAC** | **7** |

*Notes*: A special case is Val, where GTG was classed as the optimal codon for analysis based on its much higher RSCU values than its sister codons (as observed for the other optimal codons) and which had high numbers of matching tRNA genes, while its ΔRSCU was near zero; thus GTC was classed as Non-opt$_{low\text{-}tRNAs}$, but this codon may also be interpreted as optimal (wobble) status. Assuming GTC to be the optimal codon had no consequence to our results and GTT and GTA remained non-optimal in both cases. All of the optimal codons shown were classed as optimal (preferred) at the CSD (Subramanian et al. 2022), had P-values<0.05 (that used the highest 10% ENC versus the lowest 10% ENC genes in the genome (Subramanian et al. 2022)), and also strongly agreed with the G3 and C3 optimal codons obtained from comparison of the highest and least expressed genes in *D. melanogaster* (Duret and Mouchiroud 1999). RSCU values represent the values across all concatenated genes for each gene set. Herein a single optimal codon was defined per amino acid using ΔRSCU$_{Ribosome\text{-}All}$. Moreover, we classified all non-optimal codons (as all that were not the optimal codon, as described in Supplemental Text File S1), and every codon was also classified with respect to tRNA gene number status. N=169 for the ribosomal protein genes. The cutoff for classification as a high and low number of tRNA genes was ≥5 and <5 respectively (median of all tRNAs per gene was 4).

**Table S2.** The 30 developmental stages and 29 tissue types used for expression breadth analysis, or *tau*. The RPKM data is from Modencode (Li et al. 2014) and is available at FlyBase (Gramates et al. 2022). See also (Whittle and Extavour 2023). The 4d ovary and testis datasets used to assess sex-biased gene expression are included, and those used to assess 4-day virgin ovaries versus 4-day mated male accessory glands.

| Developmental stages | Tissue types |
|---|---|
| em0-2hr | A_MateF_1d_head |
| em2-4hr | A_MateF_4d_ovary |
| em4-6hr | A_MateM_1d_head |
| em6-8hr | A_VirF_1d_head |
| em8-10hr | A_VirF_4d_head |
| em10-12hr | A_MateF_20d_head |
| em12-14hr | A_MateF_4d_head |
| em14-16hr | A_MateM_20d_head |
| em16-18hr | A_MateM_4d_acc_gland |
| em18-20hr | A_MateM_4d_head |
| em20-22hr | A_MateM_4d_testis |
| em22-24hr | A_1d_carcass |
| L1 | A_1d_dig_sys |
| L2 | A_20d_carcass |
| L3_12hr | A_20d_dig_sys |
| L3_PS1-2 | A_4d_carcass |
| L3_PS3-6 | A_4d_dig_sys |
| L3_PS7-9 | P8_CNS |
| WPP | L3_CNS |
| P5 | L3_Wand_carcass |
| P6 | L3_Wand_dig_sys |
| P8 | L3_Wand_fat |
| P9-10 | L3_Wand_imag_disc |
| P15 | L3_Wand_saliv |
| AdF_Ecl_1days | A_VirF_20d_head |
| AdF_Ecl_5days | A_VirF_4d_ovary |
| AdF_Ecl_30days | WPP_fat |
| AdM_Ecl_1days | WPP_saliv |
| AdM_Ecl_5days | P8_fat |
| AdM_Ecl_30days | |

**Table S3.** Gene functional classification for all the testis-biased, ovary-biased and unbiased genes identified in *D. melanogaster*. Functions are from DAVID (Sherman et al. 2022). P-values indicate support of function from a Fisher's test. Genes could belong to more than one category.

| Gene classification | Percent | P-Value |
|---|---|---|
| **Testis-biased** | | |
| sexual reproduction | 5.2 | 4.30E-20 |
| multicellular organism reproduction | 5.6 | 1.40E-18 |
| Spermatogenesis | 1.6 | 6.20E-11 |
| sperm individualization | 2 | 6.10E-10 |
| tricarboxylic acid cycle | 1.5 | 5.80E-09 |
| sperm axoneme assembly | 1.3 | 7.60E-09 |
| mitochondrion | 7.2 | 1.10E-08 |
| cilium movement involved in cell motility | 0.9 | 3.30E-08 |
| carboxypeptidase activity | 1 | 3.80E-08 |
| sperm competition | 1.1 | 6.40E-08 |
| protein metabolic process | 0.9 | 8.80E-08 |
| motile cilium | 1.3 | 1.80E-07 |
| aminopeptidase activity | 1 | 2.30E-07 |
| protein import into mitochondrial matrix | 1.2 | 3.50E-07 |
| extracellular space | 7 | 9.30E-07 |
| **Ovary-biased** | | |
| nucleus | 45.3 | 7.90E-19 |
| protein binding | 25.2 | 8.80E-18 |
| chorion | 5.4 | 1.10E-12 |
| mRNA binding | 9.7 | 1.30E-11 |
| nucleic acid binding | 11.2 | 8.40E-09 |
| RNA binding | 13.2 | 1.20E-08 |
| cytoplasm | 32.9 | 2.90E-08 |
| mitotic cell cycle | 6.6 | 7.10E-08 |
| vitelline envelope | 2.3 | 9.10E-08 |
| polytene chromosome | 7 | 3.10E-07 |
| nucleoplasm | 8.9 | 5.60E-07 |
| oogenesis | 7.8 | 8.10E-07 |
| vitelline membrane formation involved in chorion-containing eggshell formation | 2.3 | 6.00E-06 |
| chromosome condensation | 3.5 | 1.90E-05 |
| chromatin binding | 5.8 | 2.20E-05 |
| ribonucleoprotein complex | 3.5 | 4.60E-05 |
| **Unbiased** | | |
| cytoplasmic translation | 12.9 | 3.20E-84 |
| cytosolic large ribosomal subunit | 7.3 | 1.20E-46 |
| cytosolic ribosome | 11.3 | 2.80E-44 |
| structural constituent of ribosome | 13.2 | 1.50E-37 |

| | | |
|---|---|---|
| cytosolic small ribosomal subunit | 5.7 | 1.50E-36 |
| cytosol | 28.2 | 6.30E-35 |
| ribosome | 11.5 | 2.90E-34 |
| nuclear chromosome | 5.7 | 2.50E-33 |
| proteasome complex | 4.5 | 3.00E-28 |
| protein folding | 5.6 | 1.80E-20 |
| proteasome-mediated ubiquitin-dependent protein catabolic process | 5.7 | 4.10E-19 |
| cytoplasm | 33 | 8.50E-19 |
| DNA-templated transcription, initiation | 3.7 | 4.70E-17 |
| nucleoplasm | 9.1 | 4.80E-17 |
| proteasome regulatory particle | 2.7 | 4.90E-17 |

**Table S4.** The size complexity (S/C) scores of amino acids (Dufton 1997).

| **Amino Acid** | **S/C score** |
|---|---|
| Ala | 4.76 |
| Arg | 56.34 |
| Asn | 33.72 |
| Asp | 32.72 |
| Cys | 57.16 |
| Gln | 37.48 |
| Glu | 36.48 |
| Gly | 1 |
| His | 58.7 |
| Ile | 16.04 |
| Leu | 16.04 |
| Lys | 30.14 |
| Met | 64.68 |
| Phe | 44 |
| Pro | 31.8 |
| Ser | 17.86 |
| Thr | 21.62 |
| Trp | 73 |
| Tyr | 57 |
| Val | 12.28 |

**Table S5.** The ΔRSCU$_{Testis-Ovary}$ and ΔRSCU$_{Testis-Unbiased}$ values in *D. melanogaster* for each of the 18 degenerate amino acids. The status of each codon per amino acid with respect to tRNA genes is shown. *P<0.05, **P<0.001. The primary, or most preferred, non-optimal codon per amino acid (in testis-biased genes) was defined as the codon with the largest positive ΔRSCU, which was typically the same codon identified in both ΔRSCU contrasts (in bold). In two cases, where the ranking of the largest positive codon varied between contrasts (Pro, Val) the codon with the largest average ΔRSCU was taken as the primary non-optimal codon. N/A=not applicable.

| **Amino acid** | **Codon** | **ΔRSCU$_{Testis-Ovary}$** | | **ΔRSCU$_{Testis-Unbiased}$** | | **Codon Status** |
|---|---|---|---|---|---|---|
| | | ΔRSCU | P | ΔRSCU | P | |
| Ala | GCT | +0.028 | | +0.061 | | Non-opt$_{high-tRNAs}$ |
| Ala | GCC | -0.081 | * | -0.392 | ** | Opt$_{wobble}$ |
| Ala | **GCA** | **+0.021** | | **+0.214** | ** | **Non-opt$_{low-tRNAs}$** |
| Ala | GCG | +0.0015 | | +0.087 | * | Non-opt$_{low-tRNAs}$ |
| Arg | CGT | -0.110 | | -0.266 | ** | Non-opt$_{high-tRNAs}$ |
| Arg | CGC | -0.577 | ** | -1.097 | ** | Opt$_{wobble}$ |
| Arg | CGA | +0.076 | | +0.324 | ** | Non-opt$_{high-tRNAs}$ |
| Arg | CGG | +0.067 | | +0.256 | ** | Non-opt$_{low-tRNAs}$ |
| Arg | AGA | +0.187 | ** | +0.345 | ** | Non-opt$_{low-tRNAs}$ |
| Arg | **AGG** | **+0.283** | ** | **+0.370** | ** | **Non-opt$_{low-tRNAs}$** |
| Asn | **AAT** | **+0.154** | ** | **+0.355** | ** | **Non-opt$_{low-tRNAs}$** |
| Asn | AAC | -0.175 | ** | -0.367 | ** | Opt$_{high-tRNAs}$ |
| Asp | **GAT** | **+0.043** | | **+0.191** | ** | **Non-opt$_{low-tRNAs}$** |
| Asp | GAC | -0.071 | ** | -0.208 | ** | Opt$_{high-tRNAs}$ |
| Cys | **TGT** | **+0.073** | | **+0.226** | ** | **Non-opt$_{low-tRNAs}$** |
| Cys | TGC | -0.104 | ** | -0.126 | ** | Opt$_{high-tRNAs}$ |
| Gln | **CAA** | **+0.051** | * | **+0.201** | ** | **Non-opt$_{low-tRNAs}$** |
| Gln | CAG | -0.087 | ** | -0.240 | ** | Opt$_{high-tRNAs}$ |
| Glu | **GAA** | **+0.096** | ** | **+0.232** | ** | **Non-opt$_{high-tRNAs}$** |
| Glu | GAG | -0.139 | ** | -0.262 | ** | Opt$_{high-tRNAs}$ |
| Gly | GGT | -0.034 | | +0.013 | | Non-opt$_{low-tRNAs}$ |
| Gly | GGC | -0.289 | ** | -0.450 | ** | Opt$_{high-tRNAs}$ |
| Gly | **GGA** | **+0.204** | ** | **+0.276** | ** | **Non-opt$_{high-tRNAs}$** |
| Gly | GGG | +0.115 | * | +0.170 | ** | Non-opt$_{low-tRNAs}$ |

| | | | | | | |
|---|---|---|---|---|---|---|
| His | **CAT** | **+0.064** | | **+0.159** | ** | **Non-opt$_{low\text{-}tRNAs}$** |
| His | CAC | -0.142 | ** | -0.235 | ** | Opt$_{high\text{-}tRNAs}$ |
| Ile | ATT | +0.012 | | +0.109 | ** | Non-opt$_{high\text{-}tRNAs}$ |
| Ile | ATC | -0.198 | ** | -0.437 | ** | Opt$_{wobble}$ |
| Ile | **ATA** | **+0.114** | | **+0.266** | ** | **Non-opt$_{low\text{-}tRNAs}$** |
| Leu | TTA | +0.036 | | +0.154 | ** | Non-opt$_{low\text{-}tRNAs}$ |
| Leu | **TTG** | **+0.142** | * | **+0.259** | ** | **Non-opt$_{low\text{-}tRNAs}$** |
| Leu | CTT | +0.066 | | +0.189 | ** | Non-opt$_{low\text{-}tRNAs}$ |
| Leu | CTC | -0.002 | | -0.046 | | Non-opt$_{low\text{-}tRNAs}$ |
| Leu | CTA | +0.046 | | +0.197 | ** | Non-opt$_{low\text{-}tRNAs}$ |
| Leu | CTG | -0.353 | ** | -0.818 | ** | Opt$_{high\text{-}tRNAs}$ |
| Lys | **AAA** | **+0.026** | | **+0.161** | ** | **Non-opt$_{high\text{-}tRNAs}$** |
| Lys | AAG | -0.070 | ** | -0.213 | ** | Opt$_{high\text{-}tRNAs}$ |
| Met | ATG | - | - | N/A | | |
| Phe | **TTT** | **+0.086** | | **+0.291** | ** | **Non-opt$_{low\text{-}tRNAs}$** |
| Phe | TTC | -0.120 | ** | -0.296 | ** | Opt$_{high\text{-}tRNAs}$ |
| Pro | **CCT** | **+0.090** | * | **+0.149** | * | **Non-opt$_{high\text{-}tRNAs}$** |
| Pro | CCC | -0.150 | * | -0.528 | ** | Opt$_{wobble}$ |
| Pro | CCA | 0 | | +0.235 | ** | Non-opt$_{high\text{-}tRNAs}$ |
| Pro | CCG | 0 | * | +0.135 | * | Non-opt$_{high\text{-}tRNAs}$ |
| Ser | TCT | +0.047 | * | +0.091 | * | Non-opt$_{high\text{-}tRNAs}$ |
| Ser | TCC | +0.039 | | -0.399 | ** | Opt$_{wobble}$ |
| Ser | TCA | +0.021 | | +0.176 | ** | Non-opt$_{low\text{-}tRNAs}$ |
| Ser | TCG | -0.150 | * | -0.231 | ** | Non-opt$_{low\text{-}tRNAs}$ |
| Ser | **AGT** | **+0.214** | ** | **+0.402** | ** | **Non-opt$_{low\text{-}tRNAs}$** |
| Ser | AGC | -0.196 | * | -0.064 | | Non-opt$_{high\text{-}tRNAs}$ |
| Thr | **ACT** | **+0.166** | ** | **+0.201** | ** | **Non-opt$_{high\text{-}tRNAs}$** |
| Thr | ACC | -0.037 | | -0.334 | ** | Opt$_{wobble}$ |
| Thr | ACA | -0.111 | * | +0.082 | * | Non-opt$_{high\text{-}tRNAs}$ |
| Thr | ACG | -0.051 | * | +0.042 | | Non-opt$_{low\text{-}tRNAs}$ |

| | | | | | | |
|---|---|---|---|---|---|---|
| Trp | TGG | - | - | N/A | | |
| Tyr | **TAT** | **+0.094** | | **+0.284** | ** | **Non-opt$_{\text{low-tRNAs}}$** |
| Tyr | TAC | -0.174 | ** | -0.308 | ** | Opt$_{\text{high-tRNAs}}$ |
| Val | GTT | +0.042 | | +0.157 | ** | Non-opt$_{\text{high-tRNAs}}$ |
| Val | GTC | -0.061 | | -0.140 | * | Non-opt$_{\text{low-tRNAs}}$ |
| Val | **GTA** | **+0.066** | | **+0.148** | ** | **Non-opt$_{\text{low-tRNAs}}$** |
| Val | GTG | -0.089 | ** | -0.208 | ** | Opt$_{\text{high-tRNAs}}$ |

**Table S6.** Gene functional clustering for the 212 testis-biased genes with high use of non-optimal codons in *D. melanogaster* from Fig. 2. Functions are from DAVID (Sherman et al. 2022). P-values are derived from a modified Fisher's test, where lower values indicate greater enrichment.

| **Enrichment Score: 13.43** | **No. genes** | **P-Value** |
|---|---|---|
| sexual reproduction | 25 | 3.50E-23 |
| multicellular organism reproduction | 25 | 4.40E-21 |
| extracellular space | 27 | 4.50E-11 |
| **Enrichment Score: 3.02** | | |
| behavior | 6 | 5.50E-06 |
| secreted | 9 | 2.90E-03 |
| extracellular region | 10 | 5.50E-02 |
| **Enrichment Score: 2.49** | | |
| spermatogenesis | 15 | 4.40E-06 |
| sperm axoneme assembly | 4 | 7.60E-04 |
| differentiation | 6 | 5.30E-03 |
| **Enrichment Score: 2.2** | | |
| microtubule plus-end | 4 | 3.90E-04 |
| microtubule-associated protein RP/EB | 3 | 6.30E-04 |
| protein localization to microtubule | 3 | 6.60E-04 |
| regulation of microtubule polymerization or depolymerization | 3 | 1.60E-03 |
| microtubule plus-end binding | 3 | 2.00E-03 |
| spindle midzone | 3 | 1.60E-02 |
| cytoplasmic microtubule | 3 | 1.60E-02 |
| calponin homology domain | 3 | 1.90E-02 |
| spindle assembly | 3 | 2.80E-02 |
| microtubule organizing center | 3 | 6.30E-02 |
| microtubule binding | 3 | 9.70E-02 |

**Table S7.** Examples of testis-biased genes with very high non-optimal codon use (>60%). Note Met and Trp were excluded in percentage calculations (see Supplementary Text File 1).

| FlyBase ID | Gene name | Percent non-optimal codons |
|---|---|---|
| FBgn0083938 | *BG642163* | 82.21 |
| FBgn0083936 | *Acp54A1* | 76.19 |
| FBgn0250831 | *BG642167* | 74.75 |
| FBgn0004414 | *msopa* | 74.39 |
| FBgn0011668 | *Mst57Da* | 72.97 |
| FBgn0028412 | *Mst33A* | 72.41 |
| FBgn0261055 | *Sfp26Ad* | 72.03 |
| FBgn0002855 | *Acp26Aa* | 70.77 |
| FBgn0011559 | *Acp36DE* | 70.59 |
| FBgn0011239 | *ms(2)35Ci* | 70.40 |
| FBgn0047334 | *BG642312* | 70.27 |
| FBgn0034152 | *Acp53C14a* | 69.75 |
| FBgn0051281 | *Tpl94D* | 68.15 |
| FBgn0001281 | *janB* | 67.88 |
| FBgn0015584 | *Acp53Ea* | 67.52 |
| FBgn0004175 | *Mst84Dd* | 67.14 |
| FBgn0032269 | *w-cup* | 66.33 |
| FBgn0259975 | *Sfp87B* | 66.28 |
| FBgn0004174 | *Mst84Dc* | 66.04 |
| FBgn0004173 | *Mst84Db* | 65.75 |
| FBgn0050365 | *spaw* | 65.49 |
| FBgn0036970 | *Spn77Bc* | 65.42 |
| FBgn0250832 | *Dup99B* | 64.71 |
| FBgn0020399 | *Mst89B* | 64.07 |
| FBgn0011670 | *Mst57Dc* | 63.89 |
| FBgn0000592 | *Est-6* | 63.71 |
| FBgn0013300 | *Mst35Ba* | 63.57 |
| FBgn0039124 | *tbrd-1* | 62.60 |
| FBgn0038281 | *RpL10Aa* | 62.38 |
| FBgn0067903 | *IM18* | 62.32 |
| FBgn0003742 | *tra2* | 62.07 |
| FBgn0031367 | *c-cup* | 61.34 |
| FBgn0043025 | *Msi* | 61.28 |
| FBgn0032660 | *elfless* | 61.16 |
| FBgn0046874 | *Pif1B* | 60.93 |
| FBgn0019828 | *dj* | 60.83 |
| FBgn0001099 | *gdl* | 60.64 |
| FBgn0031878 | *sip2* | 60.48 |
| FBgn0038225 | *soti* | 60.14 |

**Table S8.** The average RSA of amino acids from testis-biased genes when encoded by the primary non-optimal codon (defined in Table S5) versus the optimal codon (Table S1) for all 18 degenerate amino acids. Standard errors (SE) are shown. Sign test P<0.001.

| | Primary non-optimal codon | | | | Optimal codon | | | | |
|---|---|---|---|---|---|---|---|---|---|
| **Amino acid** | **Codon** | **Average RSA** | **SE** | **Codon status** | **Codon** | **Average RSA** | **SE** | **Codon status** | **Sign test (+/-)** |
| Ala | GCA | 0.397 | 0.003 | Non-optlow-tRNAs | GCC | 0.330 | 0.002 | Optwobble | + |
| Arg | AGG | 0.496 | 0.003 | Non-optlow-tRNAs | CGC | 0.472 | 0.002 | Optwobble | + |
| Asn | AAT | 0.540 | 0.002 | Non-optlow-tRNAs | AAC | 0.521 | 0.002 | Opthigh-tRNAs | + |
| Asp | GAT | 0.538 | 0.002 | Non-optlow-tRNAs | GAC | 0.529 | 0.002 | Opthigh-tRNAs | + |
| Cys | TGT | 0.244 | 0.003 | Non-optlow-tRNAs | TGC | 0.226 | 0.002 | Opthigh-tRNAs | + |
| Gln | CAA | 0.539 | 0.002 | Non-optlow-tRNAs | CAG | 0.516 | 0.002 | Opthigh-tRNAs | + |
| Glu | GAA | 0.577 | 0.002 | Non-opthigh-tRNAs | GAG | 0.555 | 0.001 | Opthigh-tRNAs | + |
| Gly | GGA | 0.448 | 0.003 | Non-opthigh-tRNAs | GGC | 0.426 | 0.003 | Opthigh-tRNAs | + |
| His | CAT | 0.441 | 0.003 | Non-optlow-tRNAs | CAC | 0.427 | 0.003 | Opthigh-tRNAs | + |
| Ile | ATA | 0.253 | 0.003 | Non-optlow-tRNAs | ATC | 0.219 | 0.002 | Optwobble | + |
| Leu | TTG | 0.273 | 0.003 | Non-optlow-tRNAs | CTG | 0.244 | 0.002 | Opthigh-tRNAs | + |
| Lys | AAA | 0.564 | 0.002 | Non-opthigh-tRNAs | AAG | 0.547 | 0.001 | Opthigh-tRNAs | + |
| Phe | TTT | 0.268 | 0.003 | Non-optlow-tRNAs | TTC | 0.239 | 0.002 | Opthigh-tRNAs | + |
| Pro | CCT | 0.498 | 0.003 | Non-opthigh-tRNAs | CCC | 0.455 | 0.002 | Optwobble | + |
| Ser | AGT | 0.488 | 0.003 | Non-optlow-tRNAs | TCC | 0.452 | 0.003 | Optwobble | + |
| Thr | ACT | 0.447 | 0.004 | Non-opthigh-tRNAs | ACC | 0.411 | 0.002 | Optwobble | + |
| Tyr | TAT | 0.308 | 0.003 | Non-optlow-tRNAs | TAC | 0.292 | 0.002 | Opthigh-tRNAs | + |
| Val | GTA | 0.303 | 0.004 | Non-optlow-tRNAs | GTG | 0.263 | 0.002 | Opthigh-tRNAs | + 18 of 18 amino acids positive P<0.001 |

**Table S9. RSCU values used to determine ΔRSCU for testis-biased versus ovary-biased genes.**

| Codon | Amino acid | RSCU testis (average) | RSCU ovary (average) |
|---|---|---|---|
| GCT | Ala | 0.770229 | 0.742442 |
| GCC | Ala | 1.793483 | 1.875 |
| GCA | Ala | 0.696605 | 0.675581 |
| GCG | Ala | 0.708843 | 0.707248 |
| CGT | Arg | 0.964476 | 1.074574 |
| CGC | Arg | 1.746725 | 2.32469 |
| CGA | Arg | 0.85952 | 0.783915 |
| CGG | Arg | 0.866714 | 0.799419 |
| AGA | Arg | 0.637085 | 0.449922 |
| AGG | Arg | 0.82798 | 0.545155 |
| AAT | Asn | 0.9275 | 0.773721 |
| AAC | Asn | 1.050873 | 1.226395 |
| GAT | Asp | 1.040055 | 0.996628 |
| GAC | Asp | 0.916441 | 0.987984 |
| TGT | Cys | 0.560273 | 0.487403 |
| TGC | Cys | 1.291321 | 1.396318 |
| CAA | Gln | 0.614028 | 0.562946 |
| CAG | Gln | 1.333723 | 1.421628 |
| GAA | Glu | 0.692697 | 0.596705 |
| GAG | Glu | 1.263821 | 1.403411 |
| GGT | Gly | 0.838723 | 0.873643 |
| GGC | Gly | 1.549574 | 1.838605 |
| GGA | Gly | 1.222249 | 1.018333 |
| GGG | Gly | 0.385011 | 0.269574 |
| CAT | His | 0.758493 | 0.694884 |
| CAC | His | 1.108384 | 1.250969 |
| ATT | Ile | 0.975993 | 0.963527 |
| ATC | Ile | 1.373308 | 1.572054 |
| ATA | Ile | 0.579017 | 0.465155 |
| TTA | Leu | 0.30036 | 0.26438 |
| TTG | Leu | 1.156441 | 1.014341 |
| CTT | Leu | 0.646277 | 0.580155 |
| CTC | Leu | 0.940644 | 0.942752 |
| CTA | Leu | 0.549214 | 0.50314 |
| CTG | Leu | 2.342544 | 2.696124 |
| AAA | Lys | 0.566736 | 0.54062 |
| AAG | Lys | 1.381015 | 1.451783 |
| ATG | Met | N/A | N/A |
| TTT | Phe | 0.751616 | 0.665814 |
| TTC | Phe | 1.21357 | 1.334419 |
| CCT | Pro | 0.542686 | 0.452481 |
| CCC | Pro | 1.30238 | 1.452674 |
| CCA | Pro | 0.973624 | 0.974225 |

| | | | |
|---|---|---|---|
| CCG | Pro | 1.119574 | 1.119845 |
| TCT | Ser | 0.560437 | 0.513798 |
| TCC | Ser | 1.46524 | 1.426395 |
| TCA | Ser | 0.558854 | 0.537752 |
| TCG | Ser | 1.171463 | 1.321783 |
| AGT | Ser | 0.858439 | 0.644302 |
| AGC | Ser | 1.360033 | 1.556357 |
| ACT | Thr | 0.792587 | 0.62686 |
| ACC | Thr | 1.624902 | 1.662752 |
| ACA | Thr | 0.647041 | 0.758605 |
| ACG | Thr | 0.900022 | 0.952016 |
| TGG | Trp | N/A | N/A |
| TAT | Tyr | 0.741681 | 0.647364 |
| TAC | Tyr | 1.171103 | 1.345194 |
| GTT | Val | 0.767544 | 0.725271 |
| GTC | Val | 0.970731 | 1.032171 |
| GTA | Val | 0.455197 | 0.389109 |
| GTG | Val | 1.749323 | 1.838605 |

NA=not applicable. Note that RSCU values per codon
for each amino acid are averages across all genes under study.

**Table S10. RSCU values used to determine ΔRSCU for testis-biased versus unbiased genes.**

| Codon | Amino acid | RSCU-testis (average) | RSCU unbiased (average) |
|---|---|---|---|
| GCT | Ala | 0.770229 | 0.709157 |
| GCC | Ala | 1.793483 | 2.185702 |
| GCA | Ala | 0.696605 | 0.482926 |
| GCG | Ala | 0.708843 | 0.622281 |
| CGT | Arg | 0.964476 | 1.231091 |
| CGC | Arg | 1.746725 | 2.84443 |
| CGA | Arg | 0.85952 | 0.535769 |
| CGG | Arg | 0.866714 | 0.610777 |
| AGA | Arg | 0.637085 | 0.291901 |
| AGG | Arg | 0.82798 | 0.457207 |
| AAT | Asn | 0.9275 | 0.572033 |
| AAC | Asn | 1.050873 | 1.418182 |
| GAT | Asp | 1.040055 | 0.848727 |
| GAC | Asp | 0.916441 | 1.124975 |
| TGT | Cys | 0.560273 | 0.334116 |
| TGC | Cys | 1.291321 | 1.41795 |
| CAA | Gln | 0.614028 | 0.412545 |
| CAG | Gln | 1.333723 | 1.574413 |
| GAA | Glu | 0.692697 | 0.461157 |
| GAG | Glu | 1.263821 | 1.525835 |
| GGT | Gly | 0.838723 | 0.825983 |
| GGC | Gly | 1.549574 | 2 |
| GGA | Gly | 1.222249 | 0.946116 |
| GGG | Gly | 0.385011 | 0.214992 |
| CAT | His | 0.758493 | 0.599537 |
| CAC | His | 1.108384 | 1.344314 |
| ATT | Ile | 0.975993 | 0.866727 |
| ATC | Ile | 1.373308 | 1.810678 |
| ATA | Ile | 0.579017 | 0.313322 |
| TTA | Leu | 0.30036 | 0.145901 |
| TTG | Leu | 1.156441 | 0.897388 |
| CTT | Leu | 0.646277 | 0.456975 |
| CTC | Leu | 0.940644 | 0.986727 |
| CTA | Leu | 0.549214 | 0.353554 |
| CTG | Leu | 2.342544 | 3.160926 |
| AAA | Lys | 0.566736 | 0.405851 |
| AAG | Lys | 1.381015 | 1.594298 |
| ATG | Met | N/A | N/A |
| TTT | Phe | 0.751616 | 0.460413 |
| TTC | Phe | 1.21357 | 1.510083 |
| CCT | Pro | 0.542686 | 0.393421 |
| CCC | Pro | 1.30238 | 1.830612 |
| CCA | Pro | 0.973624 | 0.738529 |

| | | | |
|---|---|---|---|
| CCG | Pro | 1.119574 | 0.984083 |
| TCT | Ser | 0.560437 | 0.46914 |
| TCC | Ser | 1.46524 | 1.864364 |
| TCA | Ser | 0.558854 | 0.383107 |
| TCG | Ser | 1.171463 | 1.402694 |
| AGT | Ser | 0.858439 | 0.456777 |
| AGC | Ser | 1.360033 | 1.424744 |
| ACT | Thr | 0.792587 | 0.591471 |
| ACC | Thr | 1.624902 | 1.95914 |
| ACA | Thr | 0.647041 | 0.564942 |
| ACG | Thr | 0.900022 | 0.857669 |
| TGG | Trp | N/A | N/A |
| TAT | Tyr | 0.741681 | 0.457471 |
| TAC | Tyr | 1.171103 | 1.479818 |
| GTT | Val | 0.767544 | 0.610198 |
| GTC | Val | 0.970731 | 1.111058 |
| GTA | Val | 0.455197 | 0.307157 |
| GTG | Val | 1.749323 | 1.958231 |

NA=not applicable. Note that RSCU values per codon for each amino acid are averages across =all genes under study.

### Supplementary Text File S1

#### Supplementary Results and Discussion

Here, we provide additional details on topics discussed in the main text Results and Discussion.

##### Identification and Characterization of the Sex-Biased Genes: Distribution of Sex-Biased Genes Across Chromosomes

The sex-biased and unbiased genes were largely similar in distribution among autosomes 2 and 3 in *D. melanogaster* (respectively, 45.0 and 46.5% of testis-biased genes, 38.7 and 45.6% of unbiased genes, and 34.5 and 36.4% of ovary-biased; less than 1% on the micro-chromosome 4). However an elevated percentage of the ovary-biased gene set was located on the X-chromosome (N=72 genes, or 27.9%) as compared to the testis-biased (N=75 or 8.2%) or unbiased genes (N=91, or 15.0%), possibly reflecting weak dosage compensation on the X chromosome in male gonads (and thus low propensity for male-bias), or innate benefits of the location of female-biased genes on the X chromosome as they are carried in females more often than autosomes (Parisi et al. 2003; Argyridou and Parsch 2018; Whittle et al. 2020).

##### Optimal and Non-optimal Codon Identified in Other *Drosophila melanogaster* subgroup species

It is worth noting that while we focused on *D. melanogaster* herein due to the extent of publicly available genomic and transcriptome datasets and gene characterization for this species, we found that four of its sister *melanogaster* subgroup species (*D. simulans, D. sechellia*, *D. yakuba* and *D. erecta*) had the exact same identity of optimal codons and non-optimal codons (Table S1), and the RSCU values and tRNA gene counts used to identify optimal/non-optimal statuses were strongly correlated to *D. melanogaster* (Spearman's R of the RSCU values within highly expressed ribosomal protein genes across all codons was $R \geq 0.990$ ($P<10^{-7}$ and for tRNA gene counts per codon was $R \geq 0.998$ ($P<10^{-7}$) suggesting the patterns we report herein for *D. melanogaster*, may also apply to its four related sister species.

##### Identification and Characterization of the Sex-Biased Genes: CAI and expression level and CDS length (Fig. S3)

We found a stepwise decrease in CAI from unbiased, to ovary-biased to testis-biased genes (Ranked ANOVA and Dunn's $P<0.05$, Fig. S3A), thus corresponding to a marked increase in non-optimal codons in the testis-biased genes. The pattern in CAI differs from prior suggestions, based on analysis solely of optimal codon use (without tRNA analysis, or non-optimal analysis in those studies), that ovary-biased and unbiased genes in *D. melanogaster* had similar Fop values (Zhang et al. 2004; Hambuch and Parsch 2005). We suggest that our results likely have greater sensitivity than previous studies, as we obtained our CAI values by using genome-wide RNA-seq herein (Graveley et al. 2011; Li et al. 2014; Gramates et al. 2022), rather than partial microarrays as done for previous studies (Zhang et al. 2004; Hambuch and Parsch 2005). Furthermore, the study of only unbiased genes that were substantially expressed within the sexual tissue types (≥100 RPKM in at least one sexual tissue (and less than 5 fold difference between sexes), see Materials and Methods) exposes a heightened tendency for optimal codon use in unbiased genes. Importantly, Fig. S3A demonstrates a marked elevation in non-optimal codon use in the testis-biased genes, as compared to ovary-biased and unbiased genes.

Given that expression level and CDS length have each been associated with optimal codon use in various organisms (Comeron et al. 1999; Duret and Mouchiroud 1999; Akashi 2001; Hambuch and Parsch 2005; Ingvarsson 2007; Whittle et al. 2007; Whittle et al. 2019; Whittle et al. 2021), and thus, indirectly linked to non-optimal codon use, we subdivided the gene sets by expression level (two classes, ≥200 RPKM, and ≥100-200 RPKM; note that all genes under study had ≥100RPKM in the tissue with sex-biased expression and thus none were classed as having low expression; the average testis and ovary RPKM was used for this assessment for unbiased genes) and into short and long CDS lengths, with long defined as ≥272 codons, which represents the lowest $33^{rd}$ percentile of the genome-wide CDS lengths. We found that the ≥200 RPKM expressed genes had an elevated CAI as compared to the ≥100-200 RPKM group for testis-biased, ovary-based and for unbiased genes (MWU-tests $P<0.05$, Fig. S3B). With respect to CDS length, short CDS genes had higher CAI than long CDS, for ovary-biased and unbiased genes (MWU-tests $P<0.05$, Fig. S3B). However, an opposite pattern was found for testis-biased genes, where the short CDS genes had markedly lower CAI than the long CDS, for both expression classes (MWU-tests $P<0.05$ for each contrast), demonstrating a unique pattern of ameliorated non-optimal codon use in short testis-biased CDS. While it may be suggested that that lower optimal codons in testis-biased genes (Fig. S3A) may be due to genetic inference (background selection or selective sweeps (Hill and Robertson 1966)) from linked adaptive amino acid substitutions causing fixation of linked non-optimal codons (Betancourt and Presgraves 2002), interference may be expected to reduce optimal codon use (and elevate non-optimal use) in longer CDS, rather than shorter CDS (Comeron et al. 1999; Betancourt and Presgraves 2002; Loewe and Charlesworth 2007), a trend opposite to that observed for testis-biased genes (Fig. 3A,B) (Hambuch and Parsch 2005). Thus, this suggests that the accumulation of non-optimal codons in testis-biased genes is influenced by factors other than genetic interference, and may reflect a different process such as selection. We therefore followed up this observation with detailed codon use analyses.

Moreover, it is notable that the extremely highly expressed testis-biased genes which were at least 5-fold sex-biased, tended to have a higher degree of sex-bias than the highly expressed testis-biased genes (the median was 149.5 and 210 for long and short genes in the former, and 71.0 and 69.5 in the latter, consistent with high extent of specialization in the testis in each group), suggesting the more-biased expression in the testis (among genes with ≥100 RPKM), the greater use of non-optimal codons

**Non-optimal Codon use is Non-random in *D. melanogaster* Sex-biased Genes (Fig. S4)**

We examined examples of non-optimal codons that had extremely high use, defined as RSCU ≥1.5, (that is, at least 1.5-fold higher than expected under equal codon use). For the Non-

$opt_{high\text{-}tRNAs}$ codons Thr-ACT and Gly-GGA, we found that 8.6% and 28.0% of the testis-biased genes had extreme use of these two respective codons, which was more than two-fold higher than that found for ovary-biased and unbiased genes (Chi-square tests P<0.05, Fig. S4A). For the Non-$opt_{low\text{-}tRNAs}$ Ile-ATA and Val-GTA, we found that more testis-biased genes had extreme use of these non-optimal codons than ovary-biased and unbiased genes. For example, nearly 4.0% of testis-biased genes had extremely high use of GTA, which was ten-fold more common than found in ovary-biased and unbiased genes (<0.4% of genes, MWU-tests P<0.05, Fig. S4A). Thus, genes with extreme use of non-optimal codons are increased in the testis-biased gene set.

It is worth noting that with respect to optimal codons, the percent of genes with extreme use of $Opt_{high\text{-}tRNAs}$ codons (RSCU ≥1.5) differed with respect to sex-biased gene sets, with examples shown in Fig. S4. Extreme use of the two $Opt_{high\text{-}tRNAs}$ codons Arg-GCG and Leu-CTG (shown in Fig. S4B) was found for 87.2% and 93.7% of the unbiased genes respectively, higher than the percentages found for ovary-biased and testis-biased genes (values between 56.1% and 89.1 %, Chi-square tests P< 0.05, Fig. S4B). A similar directional pattern was found for the $Opt_{wobble}$ codons Pro-CCC and Ile-ATC, with extreme use being more common in unbiased and ovary-biased genes than in testis-biased genes (noting that the former codon had similar percentages for unbiased and ovary-biased genes. Fig. S4B, Chi-square for all other paired tests had P<0.05).

Notably, the two optimal wobble codons had markedly fewer genes with extreme use in the unbiased gene set, than the optimal codons with high-tRNAs. For example, the $Opt_{wobble}$ codons CCC-Pro and Ile-ATC had extreme use in only 57.8% and 73.0% of unbiased genes (as compared to the 87.2% and 93.7% for $Opt_{high\text{-}tRNAs}$ Arg-GCG and Leu-CTG, Chi-square tests P<0.05), and a similar reduction in use of wobble optimal codons was found for ovary-biased and unbiased genes (Chi-square tests P<0.05, Fig. S4B). Thus, optimal wobble codons are less likely to exhibit extreme use in highly expressed sex-biased genes than optimal codons with plentiful tRNAs. Such a pattern may be a result of the fact that wobble codons have lower binding affinity to the tRNAs (Letzring et al. 2010; Stadler and Fire 2011; Quax et al. 2015; Stein and Frydman 2019), making them less efficient for translation than codons with plentiful tRNAs. In addition, it is also feasible that our findings for wobble optimal codons may indicate these codons are only used as warranted within an mRNA when required for the pacing of translation, and thus not as often used as optimal codons with plentiful tRNAs, which may ensure rapid and efficient translation (Akashi 2001; Quax et al. 2015). Overall, the patterns found here in Fig. S4B demonstrate that optimal wobble codons are not equivalent to optimal codons with plentiful tRNAs, and that the inclusion of tRNA analysis is warranted for future codon use studies, including in *D. melanogaster*, to allow these distinctions in optimal codon use.

**Amino Acids Using Non-Optimal Codons Have High Surface Exposure and Disorder: Excluding and including Trp and Met (Fig. 2)**

We compared Percent-Non-opt calculations to average RSA per gene across all amino acids. Given that Met and Typ are each encoded by one codon and were included in gene-wide RSA values, but not Percent-Non-opt calculations (which included all 18 amino acids with two or more codons), we repeated the Percent-Non-opt calculations analysis with Met and Trp included and being scored as having optimal codons given their abundant tRNAs in Table S1. Using this approach, Percent-Non-opt and Percent-Opt across genes were each nearly perfectly correlated with the values obtained when excluding these two amino acids (Spearman's $R>0.99$, $P<10^{-7}$ for all gene sets). Thus, inclusion or exclusion of Met and Trp lead to similar results (Fig. 2A-F).

**Individual Codons per Amino Acid and Direct Measurement of tRNA Abundances (referred to in Conclusions section)**

Fig. S2 suggests co-evolution of amino acid use and tRNA gene numbers (pooled across all tRNAs) in the testis-biased, ovary-biased and unbiased genes. Thus, the patterns also suggest that the testes and ovaries may have similar tRNA populations at least at the whole amino acid level. However, individual tRNAs per amino acid could in theory still vary between the gonads. Direct quantification of synonymous tRNAs per amino acid in tissues/cell types in multicellular systems remains a challenge in terms of methodology and accuracy (due to higher structure of tRNAs, high similarity in tRNA sequences; and methods used including tRNA-seq, RNA-seq, RNA-polymerase occupancy on tRNA genes and microarrays (Dittmar et al. 2006; Guimaraes et al. 2020)). However, while certain available data from mammals and *Drosophila* suggest that tRNA abundances may vary among tissues (Moriyama and Powell 1997; Plotkin et al. 2004; Dittmar et al. 2006), other data suggest that tRNA populations may be invariant among some tissues types or cells (Dittmar et al. 2006; Schmitt et al. 2014; Rudolph et al. 2016; Stein and Frydman 2019). For example, recent analysis of genes in rapidly dividing and quiescent mouse fibroblast cells showed no differences in relative abundances of individual tRNAs (using tRNA-seq), despite marked differences in the gene codon use among cell types (Guimaraes et al. 2020). In addition, prior analysis of a subset of tRNAs using microarrays (the subset of tRNAs that were assessable using a microarray approach) in testes and ovaries in humans, showed the tRNA levels were similar between the two sex organs for at least some groups of amino acids (Dittmar et al. 2006). Thus, further data will help ascertain whether tRNA populations of synonymous codons within amino acids may vary between the gonads of *D. melanogaster*.

#### Supplementary Materials and Methods

Here, we provide additional details on the Materials and Methods from the main text.

**Genome Datasets and Gonadal Expression Analyses**

For our study, we downloaded the genome of *D. melanogaster* (13,986 protein coding genes, in version 6.50 (Gramates et al. 2022)). We extracted the longest CDS per gene for study. Before the assessment of sex-biased genes, we first identified and characterized the organismal optimal and non-optimal codons in *D. melanogaster*. Sets of optimal codons, those most used in highly expressed genes have been inferred in a number of studies in *D. melanogaster,* and have been uncorrelated to intron content (Shields et al. 1988), findings consistent with selection favoring optimal codon use (Shields et al. 1988; Moriyama and Powell 1997; Powell and Moriyama 1997; Duret and Mouchiroud 1999; Heger and Ponting 2007; Payne and Alvarez-Ponce 2019) (studies typically finding GC3-ending optimal codons; sometimes multiple codons were defined as optimal per amino acid, and usually excluding non-optimal codon and tRNA analysis). To define the optimal codons for our objectives herein, we compared the RSCU of highly expressed ribosomal protein genes (N=169) to that of the entire genome (downloaded RSCU from the CSD (Subramanian et al. 2022)), denoted as $\Delta RSCU_{Ribosome-All}$. The RSCU measures the relative frequency of a codon per synonymous codon family (amino acid) as compared to that expected under random use (e.g., expected value=1 under equal use, and higher and lower values indicate greater and lower use than expected (Sharp and Li 1987)). We defined the optimal codon per amino acid as the codon with the largest positive $\Delta RSCU_{Ribosome-All}$ (shown in Table S1, each RSCU from the CSD (Subramanian et al. 2022)), which in all cases was also classed as an optimal codon (called preferred therein) at the CSD with $P<0.05$ (which used genes with high and low effective number of codons (ENC) (Subramanian et al. 2022), see Notes in Table S1). The primary optimal codon for *D. melanogaster* using this approach was largely concordant with optimal codon lists obtained using ΔRSCU of the most highly versus the most lowly expressed genes defined previously (Duret and Mouchiroud 1999). This is another stringent method to identify optimal codons (Duret and Mouchiroud 1999; Sharp et al. 2005; Cutter et al. 2006; Ingvarsson 2008; Wang et al. 2011; Whittle et al. 2011; Whittle and Extavour 2015; Whittle et al. 2019; Whittle et al. 2021), which yielded optimal codons ending with G3 and C3 (and sometimes more than one optimal codon per amino acid (Shields et al. 1988; Moriyama and Powell 1997; Powell and Moriyama 1997; Heger and Ponting 2007; Payne and Alvarez-Ponce 2019)). However, we specifically identified a single optimal codon per amino acid herein, even for amino acids encoded by three or more synonymous codons, as typically one codon was much favored over all others (see the bolded ribosomal gene RSCU values versus un-bolded per amino acid and $\Delta RSCU_{Ribosome-All}$, Table S1). In sum, a G3 or C3 optimal codon, one per amino acid, was identified from our criteria (Table S1) that had elevated use in highly expressed genes relative to the least expressed genes in the genome (Duret and Mouchiroud 1999; Subramanian et al. 2022), and relative to whole genome (Table S1).

Non-optimal codons, whose identities have been much less studied in *D. melanogaster* to date, were defined herein as all those codons per amino acid that were not the optimal codon. This classification approach was supported by the facts that first, we found that all of the non-optimal codons typically had a negative value or near-zero value for $\Delta RSCU_{Ribosome\text{-}All}$ (Table S1), and second, each non-optimal codon had a much lower absolute RSCU value in the ribosomal protein genes than the optimal codon (typically at least 1.5 fold smaller than the optimal codon, Table S1), and in some cases had a value in excess of six fold smaller than the optimal codon, thus confirming much lower use in highly expressed versus genome-wide genes (for example, for Val, RSCU for the optimal codon GTG was 1.84, but was only 0.27 for its sister non-optimal codon GTA). A total of 18 optimal and 41 non-optimal codons across 18 amino acids were identified and studied (Table S1). Further, we then classified the optimal codons and the non-optimal codons each into two groups, based on the presence of few or of plentiful exact matching tRNA gene copies in the genome, and studied their use with respect to sex-biased expression.

*tRNA Genes in the Genome per Codon*

The tRNA gene counts in an organism's genome have provided an effective proxy for the typical relative cellular tRNA abundances (Ikemura 1981; Ikemura 1985; Sharp et al. 1986; Percudani et al. 1997; Duret 2000; Akashi 2001; Cognat et al. 2008; Du et al. 2017; Whittle et al. 2019; Whittle et al. 2021). Thus, we obtained the number of genomic tRNAs per codon type for *D. melanogaster* from the GtRNA database ((http://gtrnadb.ucsc.edu) (Chan and Lowe 2016), which utilizes tRNAscan-SE to extract tRNA genes from the genome (Lowe and Chan 2016; Chan and Lowe 2019)).

*Identification of sex-biased genes*

To identify sex-biased genes, we used the modEncode database, which contains expression data (Graveley et al. 2011; Gramates et al. 2022) for a wide range of *D. melanogaster* tissue types and developmental stages (N=59 tissues/stages herein, Table S2 (Graveley et al. 2011; Li et al. 2014; Gramates et al. 2022). The entire expression dataset (Table S2) was used to measure *tau* across tissues (see below, (Yanai et al. 2005)), and thus all expression data were obtained from a single source. For the identification of sex-biased genes, we compared expression of 4-day mated testes and 4-day mated ovaries. First, from examining testes and ovaries, we identified all genes with five-fold or higher expression bias in the gonad of one sex than in the other, which is a high threshold for defining sex-bias (Proschel et al. 2006; Whittle et al. 2007; Mank et al. 2008; Meisel 2011; Assis et al. 2012). Further, for the sex-biased gene set, we only included genes that had RPKM≥100 (in the testes or the ovaries) to ensure there was high expression in at least one sex, such that a high sex-bias ratio was not obtained from two weak expression values. The cutoff of 100 RPKM was at the $90^{th}$ and $95^{th}$ percentiles of RPKM for genes from the testes and ovaries respectively (mean expression (RPKM) for the gene set per tissue type was 389.8 ± 24.3 and 349.5 ± 36.4, t-test p=0.42), affirming that highly expressed sex-biased genes were under study. Testis- and ovary-biased genes included genes that were sex-specific, or without detectable expression in the other tissue (Fig. S1). Unbiased genes were identified as those that had ≥100 RPKM in either the testes or ovaries and no sex-bias (<5-fold difference, and thus given the cutoff of 100 RPKM in one sexual tissue, had at least 20 RPKM in the opposite tissue type), thus ensuring expression in both sexes. GO analysis was conducted for testis- and ovary-biased genes in DAVID (Sherman et al. 2022) to verify the identified gene sets were associated with testis and ovaries.

Male and female reproductive expression datasets were also available from modEncode (Graveley et al. 2011; Li et al. 2014) for male accessory glands, which produce the seminal fluid proteins (Sepil et al. 2019), and for the virgin female ovaries (Table S2). Thus, for additional rigor, we assessed the fraction of the identified sex-biased genes in our main analysis of gonads of mated tissues were at least five-fold male-biased when using contrasts of the accessory glands vs virgin ovary tissues.

*Comparison of Optimal and Non-optimal Codon Use in Sex-biased Genes*

The finest scale to study codon use is on an individual codon-by-codon basis; for example, sex-differences in codon use for each of 18 optimal codons and each of the 41 non-optimal codons. For our analysis of sex-biased genes, we determined the RSCU for every codon for each gene in the testis-biased, ovary-biased and unbiased genes using the program CodonW (Peden 1999). Then, for each of the 59 codons under study, we determined $\Delta RSCU_{Testis\text{-}Ovary}$ and as $\Delta RSCU_{Testis\text{-}Unbiased}$ across all genes per gene set, as described in the main text (Whittle et al. 2007).

***Comparisons Among Sex-biased Gene Sets using Codon Use Gene Indices***

To initially screen the level of optimal codon and non-optimal codon use in the sex-biased gene sets we used the codon adaptation index (CAI) and the frequency of optimal codons (Fop) (Bahiri-Elitzur and Tuller 2021), which broadly measure codon adaptation for optimal codons, and which do not include tRNA analysis. CAI provides a score of codon use for each gene (0 to 1) based on codon frequencies in the highly expressed reference gene set (Subramanian et al. 2022), and is thought to reflect adaptation in codon use to gene expression (Sharp and Li 1987; Bahiri-Elitzur and Tuller 2021). Fop comprises an index of how often optimal codons are used in each gene under study (0 for no optimal codons used for a gene, 1 for only optimal codons) (Ikemura 1981). We found CAI and Fop values were strongly correlated across the *D. melanogaster* genome (Spearman's R=0.967, $P<10^{-7}$), and thus solely used CAI values for analysis, which were obtained from the CSD (Subramanian et al. 2022).

CAI and Fop have typically been used to assess codon adaptation in terms of optimal codon use in genes (Ikemura 1981; Sharp and Li 1987; Bahiri-Elitzur and Tuller 2021), including in *D. melanogaster* (Hambuch and Parsch 2005), and thus are indicators of improved translational efficiency (with values closer to 1). However, lower values in codon bias indices innately indicate increased non-optimal codon use, which accompanies a reduction of optimal codons (Zhou et al. 2015). Thus, herein lower values of CAI were taken as evidence of enhanced use of non-optimal codons per gene.

**Protein Folding, Relative Solvent Accessibility and Codon Use in the Sex-biased Genes**

The RSA value reflects the degree of burial or solvent exposure of an amino acid based on its local secondary and tertiary structures, or lack of well-folded structures, relative to its maximum possible surface area per amino acid (in Angstroms) (Braun 1998; Tien et al. 2013). To determine the RSA per amino acid we used NetsurfP v3.0 at default parameters (Hoie et al. 2022). In addition to RSA, we also determined the type of secondary structure that each amino acid was a part of, that is, an alpha-helix, string (β-string/sheet) or a random coil (typically disordered) using NetsurfP 3.0 (Hoie et al. 2022).

Protein folding predictions for *D. melanogaster* proteins (Uniprot proteome Identity UP000000803, taxon identity = 7227) were determined using Alphafold2 (Juniper et al. 2021), and are available from the AlphaFold2 database for most proteins (https://alphafold.ebi.ac.uk)(Varadi et al. 2022). We recognize that the accuracy of predicted formations for disordered regions may sometimes be less well statistically supported than for well-structured regions in the database (Baek and Kepp 2022; Varadi et al. 2022).

**Expression Specificity, or *tau***

While gene expression level has been associated with the use of optimal codons, and with translational efficiency of abundant transcripts (Ikemura 1985; Shields et al. 1988; Akashi 2001; Hambuch and Parsch 2005; Behura and Severson 2011; Wang et al. 2011; Galtier et al. 2018; Whittle et al. 2019; Whittle et al. 2021), a factor worth consideration herein with a possible effect on codon use is expression breadth. Thus, we used data on the expression specificity index *tau* for each studied gene in the *D. melanogaster* genome using gene expression data available across the 59 tissues/stages from the modEncode database (Table S2 (Graveley et al. 2011; Gramates et al. 2022) as per (Yanai et al. 2005) (see also our prior assessment of *tau* in (Whittle and Extavour 2023)). *tau* values can range from 0 to 1, where values nearer to 1 indicate high expression specificity and lower values denote greater expression breadth across a range of tissue types (Yanai et al. 2005; Whittle and Extavour 2023). As shown in Table S2, the data included gene expression levels (RPKM) across development for the embryos (12 stages), larvae (6 stages), pupae (6 stages) and adults (3 stages of males/females), and for tissue types of the adult males and females. For each gene, the *tau* value was measured as follows:

$$\textit{tau}\ (\tau) = \sum_{i=1}^{n}(1-\hat{x}_i)/(n-1);\ \ \hat{x}_i = x_i/max\ (x_i)$$

where $n$=number of tissues/stages, $i$=tissue/stage, $x_i$= transcription level of gene in tissue/stage $i$, and max ($x_i$)= the transcription level in the tissue/stage type with maximum expression (Yanai et al. 2005).
